# Atg1 phosphorylates Atg2 to gate phospholipid flow through a continuous conduit for autophagosome biogenesis

**DOI:** 10.1101/2025.05.24.655882

**Authors:** Yuji Sakai, Tetsuya Kotani, Tatsuro Maruyama, Kazuaki Matoba, Li Hao, Kuninori Suzuki, Chika Kakuta, Junko Shimasaki, Yuji Sugita, Tatsuya Niwa, Yuko Fujioka, Takuo Osawa, Hitoshi Nakatogawa, Nobuo N. Noda

## Abstract

Autophagosome biogenesis requires delivery of millions of phospholipids. In budding yeast, Atg2 and Atg18 couple the ER to the isolation membrane for lipid transport, yet the mechanism is poorly understood. Combining molecular dynamics simulations, in vitro imaging-based lipid transfer assays, and in vivo analyses, we show that Atg1-mediated phosphorylation activates Atg2 as a lipid-transfer bridge. Simulations reveal that Atg2 contains a water-excluding hydrophobic cavity that accommodates ∼25 phospholipids with bilayer-like fluidity. Atg1 phosphorylates an N-terminal helix of Atg2 in vitro and in vivo, thereby opening the ER-facing entrance to the Atg2 cavity and enabling lipid flow between membranes in silico and in vitro. Consistently, phosphodeficient but not phosphomimetic mutations impair autophagy. Cavity narrowing along the conduit disrupts lipid alignment in silico, phosphorylation-activated bridge-type lipid transfer in vitro, and ER-to-isolation-membrane lipophilic dye flow and autophagosome expansion in vivo. These data suggest that Atg1 gates Atg2-mediated bridge-type lipid transport to build autophagosomes.

## Main text

Macroautophagy (hereafter autophagy) is an intracellular degradation system that helps maintain cellular homeostasis by breaking down various cytoplasmic components, such as proteins, organelles, and even invading bacteria, thereby contributing to the suppression of diverse diseases^1,2^. The most remarkable event in autophagy is the de novo generation of a double-membrane organelle termed the autophagosome, which sequesters biomolecules and organelles in its lumen and delivers them to the lysosome for degradation^3^. This structure expands rapidly: a single autophagosome contains millions of phospholipids, and in budding yeast its formation is completed on the order of minutes^4,5^. Genetic, imaging, and lipidomic studies have implicated multiple membrane sources, with the endoplasmic reticulum (ER) emerging as a major contributor^6–9^. Yet the central mechanistic question remains unresolved: how bulk phospholipids are mobilized from the ER and delivered to the growing isolation membrane (IM; phagophore) with sufficient speed, directionality, and spatial precision.

In *Saccharomyces cerevisiae*, this problem is addressed by core Atg proteins that assemble at the pre-autophagosomal structure (PAS) by phase separation and organize ER–IM contact sites^10–12^. Atg2, the largest core Atg protein, forms a stable complex with the phosphoinositide-binding protein Atg18. The complex is recruited to the IM in part through Atg18 recognition of phosphatidylinositol 3-phosphate (PI3P), and Atg2 also engages the ER, thereby tethering the two membranes^11–15^. Assembly of the Atg9–Atg2–Atg18 complex confines Atg2 to IM rims and helps establish functional ER–IM contact sites^16–18^. ER exit sites (ERES) were identified as ER subdomains that form contacts with the IM^11,12^, and TRAPPIII- and Ypt1-dependent establishment of these ERES–IM contact sites promotes local PI3P production and Atg18 recruitment, thereby coupling contact-site assembly to IM expansion^19^. In mammals, ATG2A has also been reported to function at ER–IM contacts^20^, and depalmitoylation of ATG2A is required for this process^21^. Biochemical and structural studies then established that Atg2 and mammalian ATG2A/B can transfer phospholipids between membranes in vitro^20,22,23^. Together with the discovery that Atg9/ATG9A acts as a phospholipid scramblase, these findings led to an appealing model in which Atg2 transfers lipids from the cytosolic leaflet of the ER to that of the IM, while Atg9 distributes newly supplied lipids between the two leaflets to drive membrane expansion^4,24–27^. Still, the exact contribution of Atg2-mediated lipid transfer to autophagosome assembly remains uncertain, as vesicular transport has also been proposed to participate^28–31^.

A major challenge is distinguishing how Atg2 moves lipids. Classical lipid-transfer proteins typically shield one or a few hydrophobic ligands inside a pocket and shuttle between membranes^32^. By contrast, Atg2 proteins belong to an emerging class of large bridge-like lipid transfer proteins (BLTPs) whose extended hydrophobic cavities or grooves could span membrane-contact-site distances and allow many glycerophospholipids to flow between bilayers^20,22,23,33–36^. Coarse-grained MD simulations further predicted that numerous phospholipids can bind within and populate these extended cavities^36–38^. The bridge model is attractive because it could support the high flux needed for organelle biogenesis, but it has been difficult to test rigorously. Conventional ensemble fluorescence resonance energy transfer (FRET)-based lipid-transfer (LT) assays report lipid dilution between vesicle populations and therefore cannot unambiguously distinguish continuous bridge flow from shuttle-like transfer, transient lipid extraction into a protein cavity, or dequenching caused by membrane tethering^20,34^. Similarly, previous hydrophobic-to-charged substitutions in lipid-binding cavities have provided important evidence that hydrophobic pathways are required^20,39–41^, but charged mutations could also impair lipid extraction or shuttle-type carrier activity. A more stringent bridge-specific test would physically constrict the pathway at an internal position without altering the amino acid composition of the hydrophobic cavity lining or its entrances: only a mechanism requiring lipids to traverse the full-length conduit would be predicted to be highly sensitive to such a constriction.

Atg2 regulation poses a second unresolved question. Autophagosome formation is initiated by the Atg1 kinase complex, which organizes the PAS by phase separation and phosphorylates multiple Atg factors^42,43^. Atg2 has been reported as a phosphorylation target^44^, but the functional consequences of this phosphorylation have only recently begun to emerge. Recent studies showed that Atg1 phosphorylation of a phospho-FFAT motif in Atg2 promotes its interaction with the ER-resident VAP protein Scs2 and, together with the Atg2 N-terminal region, provides spatiotemporal control of Atg2 association with the ER^45,46^. In parallel, Achleitner et al. proposed that Scs2 recruits the Atg1 complex to the ER through Atg13 and subsequently binds phosphorylated Atg2, thereby positioning Atg2 at ER–Atg9 vesicle contact sites^47^. These findings explain how Atg2 is recruited to the correct ER–IM contact site, but leave open a critical mechanistic question: whether Atg1 directly gates lipid entry into, or transport through, the Atg2 bridge. Such regulation would be conceptually important: if Atg2 functions as a bulk lipid conduit, its activity must be activated only at the correct membrane contact site to avoid uncontrolled lipid flow from the ER.

Here, we combine all-atom and coarse-grained MD simulations, in vivo analyses of ER-to-IM lipophilic-dye flow and autophagosome expansion, and a GUV–SLB reconstitution assay that preferentially detects bridge-type lipid transfer. We show that Atg2 forms a continuous, water-excluding conduit that accommodates ∼25 phospholipids in a fluid single file. Atg1 phosphorylates the N-terminal H2 helix in vitro and in vivo, and simulations predict that this modification opens the ER-facing entrance to enable donor-side lipid loading and intermembrane transfer. Critically, physically constricting the cavity without altering its hydrophobic lining or entrances disrupts lipid passage in silico, ER-to-IM dye flow and IM expansion in cells, and Atg1-activated bridge transport in vitro, while preserving activity in a conventional FRET-based assay. These findings demonstrate that autophagosome biogenesis requires lipid passage through an uninterrupted Atg2 conduit and identify Atg1 phosphorylation as a spatial gate that activates bridge-type lipid transport at ER–IM contact sites.

## Results

### Architecture of the Atg2–Atg18 complex

AlphaFold2 prediction of *S. cerevisiae* Atg2 yielded a ∼210 Å rod composed primarily of β-sheets, with a ∼140 Å hydrophobic cavity extending from an N-terminal region (NR; residues 1–228), through five repeat regions (RR1–RR5; residues 289–907), to a C-terminal region (CR; residues 1026–1041 and 1129–1346) (Fig. 1a)^48,49^. The seven cavity-forming units create a continuous path with a radius of ∼5–6 Å^50^ and a spiral lateral slit (Fig. 1b, 1c; Supplementary Fig. 1a). Its dimensions match the lipid-bound crystal structure of the *Schizosaccharomyces pombe* Atg2 N terminus^22^. Recent cryo-EM analysis of the yeast Atg2–Atg18 complex confirmed the tube-like architecture of the resolved central and C-terminal portions, and visualized diffuse lipid density along the resolved internal cavity^38^. Thus, the predicted full-length model provides an experimentally grounded framework for atomic-level analysis of the complete conduit.

**Figure 1.**
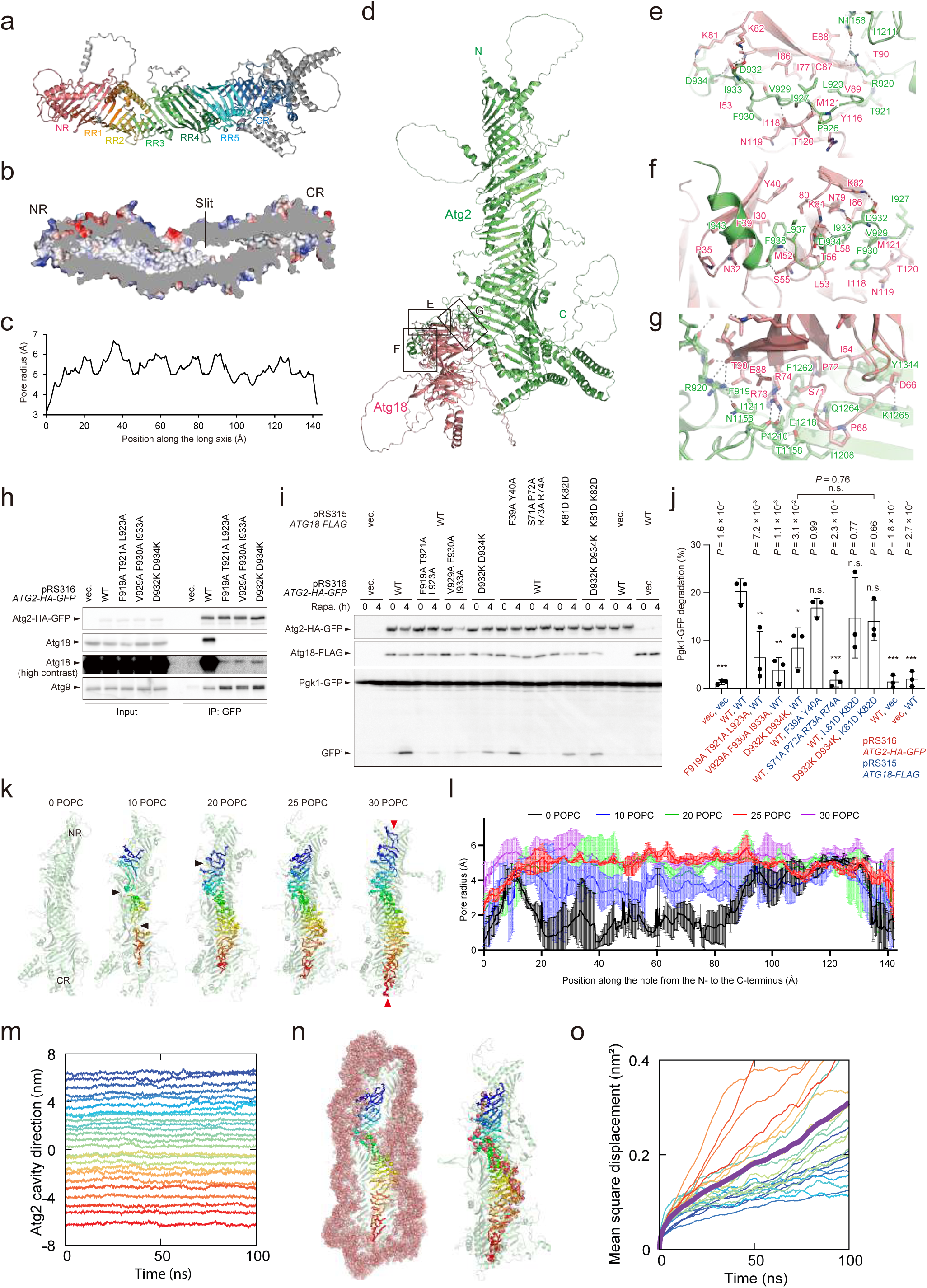
Architecture and lipid binding of the Atg2–Atg18 complex. **a,** AlphaFold model of Atg2. **b,** Cross section of the hydrophobic cavity of the rod region of Atg2. Coloring is based on the electrostatic potential (blue and red represent positive and negative electrostatic potentials, respectively). **c,** Pore radius of the Atg2 cavity from NR to CR calculated using Coot^50^. **d,** AlphaFold model of the Atg2–Atg18 complex. **e–g,** Close-up view of the interactions between Atg2 and Atg18 predicted by AlphaFold. Possible hydrophilic interactions between Atg2 and Atg18 are shown with broken lines. **h,** Coimmunoprecipitation experiments using anti-GFP nanobody-conjugated beads and cells expressing Atg2 mutants. **i,** Measurement of bulk autophagy activity by the Pgk1–GFP processing assay. Atg2 and Atg18 levels, and Pgk1–GFP degradation were examined by immunoblotting using antibodies against HA, FLAG, and GFP, respectively. **j,** Graphs show mean ± s.d. (n = 3) of ratio GFP′/(Pgk1–GFP + GFP′) in (i). *P < 0.05; ***P < 0.001; ****P < 0.0001; n.s., not significant (Tukey’s multiple comparisons test, n = 3 independent experiments). **k,** POPC distribution in the hydrophobic cavity of Atg2 after 100-ns all-atom MD simulations. Black arrowheads indicate where POPC interactions are interrupted, whereas red arrowheads indicate POPC molecules exiting the cavity. **l,** Pore radius of the Atg2 cavity after a 100-ns MD simulation. Error bars indicate s.d. of three independent calculations. **m,** Time evolution of the intermolecular distances between POPC molecules along the Atg2 cavity during the final 100 ns of 200-ns MD simulations. **n,** Water distribution. Water molecules are shown as red spheres. Left, all water molecules in the vicinity of Atg2; right, only those in contact with lipids within the Atg2 cavity. All contacting water molecules accessed the hydrophilic lipid headgroups through the cavity slit, whereas the lipid acyl chains within the cavity were shielded from water. **o,** Mean-square displacement of 25 POPC molecules in the Atg2 cavity during the final 100 ns of 200-ns MD simulations.

AlphaFold2-Multimer^49,51^ placed the Atg2 loop spanning residues 908–1025 around blades 1–3 of Atg18, which is reminiscent of the WIPI-interacting region (WIR) motif in human ATG2A binding to WIPI3^52^, with additional contacts between the Atg2 rod and blade 2 (Fig. 1d–1g). This arrangement is consistent with reports that blade 2 of Atg18 is responsible for Atg2 binding^53,54^ and agrees with the cryo-EM structure, including the principal Atg18-binding interface around Atg2 residues 922–934, although some local contacts differ (Supplementary Fig. 1b)^38^. We designed point mutants of the predicted interacting residues in Atg2 and Atg18 and evaluated protein interactions by coimmunoprecipitation and autophagic activity by monitoring Pgk1–GFP processing. Substitutions in the Atg2 loop (Phe919, Thr921, Leu923, Val929, Phe930, Asp932, Ile933, and Asp934) reduced Atg18 binding and autophagic flux, while most preserved Atg9 binding (Fig. 1h–1j). Conversely, simultaneous substitution of Ser71, Pro72, Arg73, and Arg74 in Atg18 with Ala disrupted binding to Atg2 and strongly impaired autophagy despite normal protein expression (Fig. 1i–1j; Supplementary Fig. 1c).

These data validate the Atg2–Atg18 model used below while shifting the central mechanistic question to lipid organization, conduit continuity, and donor-side gating.

### Phospholipids form a fluid, water-excluding single file within Atg2

To determine how the cavity accommodates phospholipids, we placed 0, 10, 20, 25, or 30 POPC molecules at regular intervals, equilibrated the systems by coarse-grained MD, and then performed unrestrained all-atom MD (Fig. 1k). Without lipid, the cavity collapsed to <1 Å in radius; 10 POPC molecules prevented complete collapse but formed discontinuous tail-to-tail clusters. By contrast, 25 POPC molecules formed a stable side-by-side array from the NR to the CR, with headgroups facing the lateral slit and the cavity maintained at 5–6 Å (Fig. 1k–1l). Previous coarse-grained MD studies modeled the Atg2 cavity with 15 lipids or deliberately oversaturated it with 42 lipids^37,38^. By explicitly evaluating multiple occupancy states using all-atom MD, our analysis provides a systematic estimate of the cavity’s lipid capacity under all-atom conditions. Consistent with this model, cryo-EM analysis of native *C. elegans* LPD-3 revealed ordered phospholipids arranged in a continuous row along its tunnel, with a packing density similar to that of a lipid bilayer. However, the wider, variable-diameter LPD-3 tunnel accommodates multiple lipid rows and was estimated to hold ∼72 lipids^33^. Similarly, cryo-EM analysis of human VPS13A revealed lipid-like density, including rows of fatty acyl chains, within its hydrophobic bridge domain^36^. During MD simulations, lipids rarely overtook one another (Fig. 1m), whereas loading 30 POPC molecules caused spontaneous ejection from the cavity (Fig. 1k). The 25-POPC state excluded water from the hydrophobic core and supported lipid diffusion at 1.3 μm² s ¹, comparable to lateral diffusion in bilayers^55^ (Fig. 1n, 1o). Thus, Atg2 can hold ∼25 phospholipids as a fluid, water-shielded single file rather than as tightly bound ligands.

### Atg18–PI3P engagement enables acceptor-side release but not donor-side loading

We next examined how the Atg2–Atg18 complex bridges membranes and mediates lipid transfer. We used the full-length AlphaFold2 model loaded with 28 POPC molecules, the maximum number that did not cause spontaneous lipid ejection from the cavity. Guided by PPM server predictions (Supplementary Fig. 2a)^56^, we first placed the C terminus of the complex on a planar PI3P-containing membrane resembling the IM. During a 200-ns all-atom MD simulation, Atg2–Atg18 maintained its inter-subunit interactions and remained stably associated with the membrane, positioning the cavity exit near the bilayer surface (Fig. 2a; Supplementary Fig. 2b; Supplementary Video 1). The complex induced positive membrane curvature with a radius of approximately 25 nm at the contact site and further lifted the bilayer locally toward the cavity exit (Fig. 2a–2c). A crescent-shaped region in the CR containing four amphipathic helices may act analogously to BAR domains^57^ to sense or generate curvature resembling that of the IM rim^58,59^. The FRRG motif of Atg18 engaged PI3P and contributed to local membrane lifting and bending^60^ (Fig. 2c; Supplementary Fig. 2c; Supplementary Video 1). As the cavity exit approached the membrane, a POPC molecule spontaneously entered the bilayer, and the adjacent lipid moved into the vacated position (Fig. 2a, 2c; Supplementary Fig. 2d). Neither membrane curvature nor lipid transfer occurred with a POPC-only membrane or when the PI3P-containing membrane was constrained to remain planar (Fig. 2d; Supplementary Fig. 2e, 2f), indicating that Atg18–PI3P-dependent membrane deformation is important for lipid release from the cavity, which is consistent with recent coarse-grained MD simulations^38^.

**Figure 2.**
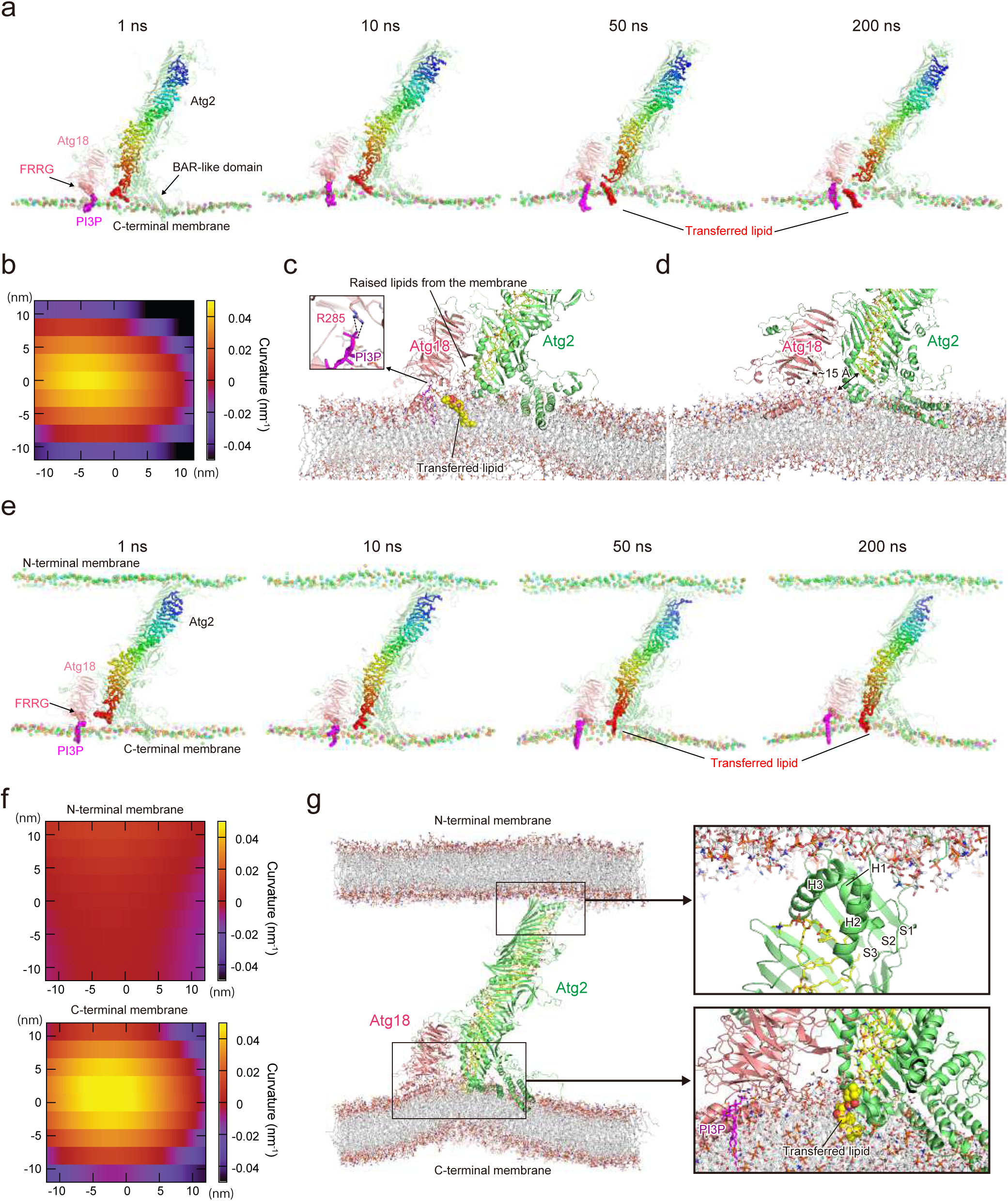
MD simulation of membrane bridging and lipid transfer by the Atg2–Atg18 complex. **a,** All-atom MD simulation of the Atg2–Atg18 complex bound to a membrane with an autophagic lipid composition. The spheres indicate phosphorus atoms in the membrane leaflet interacting with Atg2. **b,** Local curvature of the membrane averaged over the 100–200-ns interval of a 200-ns MD simulation. **c,** Close-up view of the interactions between the Atg2–Atg18 complex and the membrane with an autophagic lipid composition after a 200-ns MD simulation. **d,** Close-up view of the interactions between the Atg2–Atg18 complex and the POPC-only membrane after a 200-ns MD simulation. **e,** All-atom MD simulation of the Atg2–Atg18 complex bound to two membranes. The spheres indicate phosphorus atoms in the membrane leaflet interacting with Atg2. **f,** Local curvature of the N- and C-membranes averaged over the 100–200-ns interval of a 200-ns MD simulation. **g,** Close-up view of the interactions between the Atg2–Atg18 complex and the two membranes after a 200-ns MD simulation.

We next performed an all-atom MD simulation in which the N and C termini of Atg2–Atg18 contacted separate planar membranes: an ER-like N-membrane and a PI3P-containing IM-like C-membrane. The complex remained associated with both membranes throughout the 200-ns simulation (Fig. 2e–2g; Supplementary Video 2). Interaction with the N-membrane was mediated by hydrophobic residues in the N-terminal tail (Met1, Ala2, Phe3, Trp4, Leu5, and Pro6) and adjacent loops (Val34, Ile36, Tyr261, Met262, and Met265), which inserted into the bilayer (Supplementary Fig. 2g). The C-membrane developed positive curvature and was locally lifted toward the cavity exit, allowing POPC transfer from the cavity into the membrane, whereas the N-membrane remained planar and no lipid movement occurred between the membrane and the N-terminal cavity (Fig. 2e–2g). Coarse-grained MD simulations using 30-nm vesicles reproduced this asymmetry: lipid transfer from the C terminus to the IM-like membrane occurred within 100 ns, whereas transfer at the N terminus was not detected either with a single N-terminal vesicle or with the complex bridging two vesicles (Supplementary Fig. 2h). A recent coarse-grained MD study reported N-terminal lipid movement, but its simulation loaded 42 lipids into the Atg2 cavity, well above the ∼25-lipid capacity suggested by our all-atom MD simulations^38^. Together, these results suggest that Atg2–Atg18 bridges two membranes and promotes lipid release into the PI3P-rich C-terminal membrane, whereas lipid loading from the N-terminal membrane requires an additional activating mechanism.

### Phosphorylation of Helix 2 enables Atg2 to transfer lipids via the bridge model in silico

Upon careful inspection of the Atg2 structure following the MD simulations, we found that the extreme N-terminal region of Atg2 comprised three helices (H1–H3) and three β-strands (S1–S3) and adopted a conformation in which the first and second helices, H1 and H2, occluded the entrance to the cavity, thereby preventing phospholipid movement between the N-membrane and the cavity (Fig. 2g). This observation suggested that displacement of H1 and H2 would be required to open the cavity entrance. A recent preprint reported that point mutations in the helix corresponding to H2 of human ATG2A induced a conformational change that enabled lipid entry into the cavity and increased in vitro LT activity^61^. We therefore hypothesized that a mechanism may exist in cells to induce conformational changes in these N-terminal helices. In AlphaFold predictions, these helices, particularly H2, exhibit low pLDDT scores (Supplementary Fig. 3a), which have been associated with locally flexible or conformationally heterogeneous regions^62^. Protein phosphorylation is a common mechanism for relieving autoinhibition by disrupting or remodeling intramolecular interactions^63^. We therefore decided to examine the possibility that the N-terminal region of Atg2 is also subject to phosphorylation-dependent conformational regulation.

Atg2^H1^ does not contain any Ser/Thr residues, whereas Atg2^H2^ contains Ser109, Thr114, and Ser116 (Supplementary Fig. 3b). We therefore focused on Atg2^H2^ and used AlphaFold3 to analyze the structural consequences of introducing phosphate groups at these three Ser/Thr residues^64^. AlphaFold3 predicted displacement of Atg2^H2^ from the main body of Atg2, resulting in an open entrance to the cavity (Supplementary Fig. 3c). These results suggest that phosphorylation of the three Ser/Thr residues in Atg2^H2^ would open the entrance to the cavity and enable lipid movement between the membrane and the cavity.

We then positioned the N- and C-membranes against the predicted structure of the Atg2–Atg18 complex phosphorylated at these three residues and performed all-atom MD simulations (Fig. 3a–3e; Supplementary Video 3). As with nonphosphorylated Atg2, no curvature was induced in the N-membrane (Supplementary Fig. 3d, top). However, upon displacement of H2, H1 rotated by approximately 50°, opening the entrance to the cavity and inducing the formation of a β-strand (S0) (Fig. 3c, 3d). This created a new cavity, with S0–S3 forming the floor and H1 and H3 forming the two side walls. Furthermore, in phosphorylated Atg2, H1 inserted more deeply into the N-membrane than in unphosphorylated Atg2. H3, the S0–S1 loop, and the S2–S3 loop, specifically, residues Val34, Ile36, Lys39, Thr74, Val75, Pro153, Asn156, Tyr159, Thr160, Leu161, Met164, Lys167, Val171, and Lys175, directly interacted with the N-membrane through extensive hydrophobic and additional hydrophilic interactions, bringing the Atg2 cavity into close apposition with the membrane (Supplementary Fig. 3e). Four lipid molecules entered the newly formed cavity from the N-membrane while maintaining continuous contact with the membrane; concurrently, lipid molecules moved from the C-terminal end of the cavity into the C-membrane (Fig. 3a, 3c; Supplementary Video 3). As a result, the N- and C-membranes became connected by a single-file array of lipids within the cavity (Fig. 3e). Together, these observations suggest that H2 phosphorylation opens and docks the N-terminal entrance of the Atg2 cavity to the N-membrane, establishing continuous lipid connectivity between the N-membrane, the Atg2 cavity, and the C-membrane and thereby enabling bridge-type lipid transfer by Atg2–Atg18.

**Figure 3.**
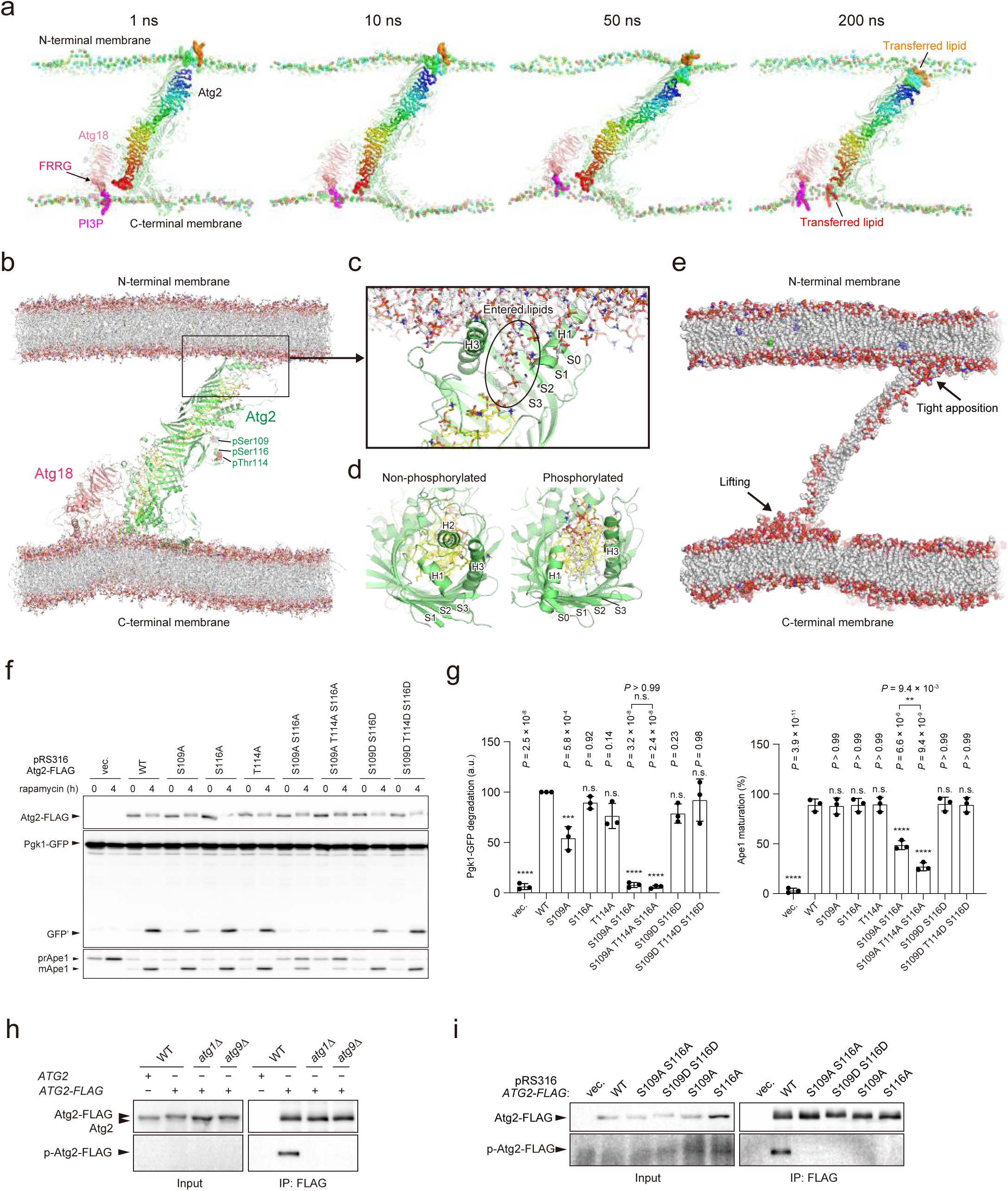
Atg1-mediated phosphorylation of Atg2^H2^ enables bridge-type LT in silico and supports autophagy in vivo. **a,** All-atom MD simulation of the phosphorylated Atg2–Atg18 complex bound to two membranes. The spheres indicate phosphorus atoms in the membrane leaflet interacting with Atg2. **b,** Close-up view of the interactions between the phosphorylated Atg2–Atg18 complex and the two membranes after a 200-ns MD simulation. **c,** Close-up view of the interactions between the N-terminal region of phosphorylated Atg2 and the N-membrane after a 200-ns MD simulation. **d,** Structural comparison of the N-terminal regions of nonphosphorylated and phosphorylated Atg2 after 200-ns MD simulations. **e,** Distribution of lipid molecules after a 200-ns MD simulation of the phosphorylated Atg2–Atg18 complex. **f,** Measurement of bulk autophagy activity by the Pgk1–GFP processing assay and Ape1 maturation. Atg2 level, Pgk1–GFP degradation, and Ape1 maturation were examined by immunoblotting using antibodies against FLAG, GFP, and Ape1, respectively. **g,** Graphs show mean ± s.d. (n = 3) of ratio GFP′/(Pgk1–GFP + GFP′) and ratio mApe1/(mApe1 + prApe1) in (f). ***P < 0.001; ****P < 0.0001; n.s., not significant (Tukey’s multiple comparisons test, n = 3 independent experiments). **h,** Coimmunoprecipitation experiments using anti-FLAG antibody-conjugated beads and cells expressing Atg2–FLAG. **i,** Coimmunoprecipitation experiments using anti-FLAG antibody-conjugated beads and cells expressing Atg2 mutants.

### Atg1-mediated phosphorylation of Atg2^H2^ in vivo is important for autophagy

In parallel with in silico studies, we examined whether the Ser/Thr residues in Atg2^H2^ are important for autophagy by monitoring Pgk1–GFP processing and Ape1 maturation (Fig. 3f, 3g). Individual substitution of Thr114 or Ser116 with Ala had little effect on autophagic activity, whereas substitution of Ser109 with Ala partially but significantly reduced autophagic activity in the Pgk1–GFP processing assay. Simultaneous substitution of Ser109 and Ser116 with Ala severely reduced autophagic activity, and substitution of all three residues with Ala caused an even more severe reduction in the case of Ape1 maturation. By contrast, substitution with Asp did not reduce autophagic activity. AlphaFold3 predictions indicated that substitution of Ser109, Thr114, and Ser116 with Ala did not alter the H2 structure, whereas substitution of all three residues with Asp, which often mimics phosphorylation, disrupted the helical structure (Supplementary Fig. 3f). These results suggest that phosphorylation of Ser109, Thr114, and Ser116 is important for autophagic activity.

The kinase activity of Atg1 is known to be important for the progression of autophagosome formation^65^. Upon starvation, the Atg1 complex undergoes phase separation to organize the PAS, where Atg1 is activated by autophosphorylation and phosphorylates various Atg factors, including Atg2^10,44^. However, there have been no reports to date showing that Atg2^H2^ is phosphorylated in vivo by Atg1 or other kinases. Therefore, we examined whether Atg2^H2^ can indeed be phosphorylated by Atg1 in vivo. We recently reported a method for inducing Atg2 phosphorylation by Atg1 in yeast: Atg1 is fused to a Gcn4 coiled-coil domain for constitutive activation and to a GFP nanobody, and this construct is co-expressed with Atg2–mGFP^45^. This allows constitutively active Atg1 to bind to mGFP-tagged Atg2 and induce its phosphorylation in an autophagy-independent manner. Atg2 was then immunoprecipitated, and phosphorylation sites were identified by LC-MS/MS (Supplementary Data 1). Several dozen phosphorylation sites were identified, including Ser109, Thr114, and Ser116 at Atg2^H2^. To determine whether Atg2^H2^ is phosphorylated under non-artificial conditions, we generated an antibody against Atg2^H2^ phosphorylated at Ser109 and Ser116 (anti-Ser109/116-P) (Supplementary Fig. 3g). Atg2–3xFLAG was immunoprecipitated with an anti-FLAG antibody from yeast expressing Atg2–3xFLAG under rapamycin treatment and analyzed by Western blotting using the anti-Ser109/116-P antibody. Phosphorylated Atg2 was detected in wild-type yeast cells, but not in atg1Δ or atg9Δ yeast cells (Fig. 3h). Both single and double Ala or Asp substitutions at Ser109 and Ser116 abolished the phosphorylated Atg2 band, confirming that the antibody specifically recognizes phosphorylation of these serine residues (Fig. 3i). Considering that Atg9 is required for PAS localization of Atg2 but not for Atg1 kinase activation, these results suggest that Atg2^H2^ is phosphorylated in vivo and that this phosphorylation is mediated by Atg1 at the PAS.

### In vitro phosphorylation of Atg2^H2^ by Atg1 enhances the FRET-based lipid transfer activity of Atg2

Next, we examined in vitro whether phosphorylation by Atg1 promotes the lipid transfer activity of Atg2. When purified Atg2 was phosphorylated with purified Atg1, Atg2^WT^ was detected with the anti-Ser109/116-P antibody (Fig. 4a). In contrast, the signal was strongly reduced in Atg2^S109A/S116A^, suggesting that Atg1 phosphorylates Ser109 and Ser116 of Atg2 in vitro. We then analyzed the effect of Atg1-mediated phosphorylation on the lipid transfer activity of Atg2 using a conventional FRET-based lipid transfer (LT) assay (Fig. 4b). Donor liposomes containing NBD-PE and Rhod-PE were mixed with unlabeled, PI3P-containing acceptor liposomes. Upon addition of Atg2–Atg18, transfer of fluorescent lipids between the two liposome populations by Atg2 dilutes the FRET pair, thereby reducing NBD–Rhod FRET and increasing NBD fluorescence. We then analyzed how the presence or absence of Atg1 and ATP affected LT activity. For Atg2^WT^, LT activity was markedly increased only when both Atg1 and ATP were present (Fig. 4c). In contrast, such an increase in activity was not observed with Atg2^S109A/S116A^ (Fig. 4d). Atg2^S109D/S116D^ exhibited higher activity than Atg2^WT^ without phosphorylation by Atg1, and Atg1 caused only a modest additional increase in activity (Fig. 4e). These results are consistent with those recently reported in a preprint^47^ and strongly suggest that phosphorylation of Atg2^H2^ by Atg1 enhances its lipid transfer activity.

**Figure 4.**
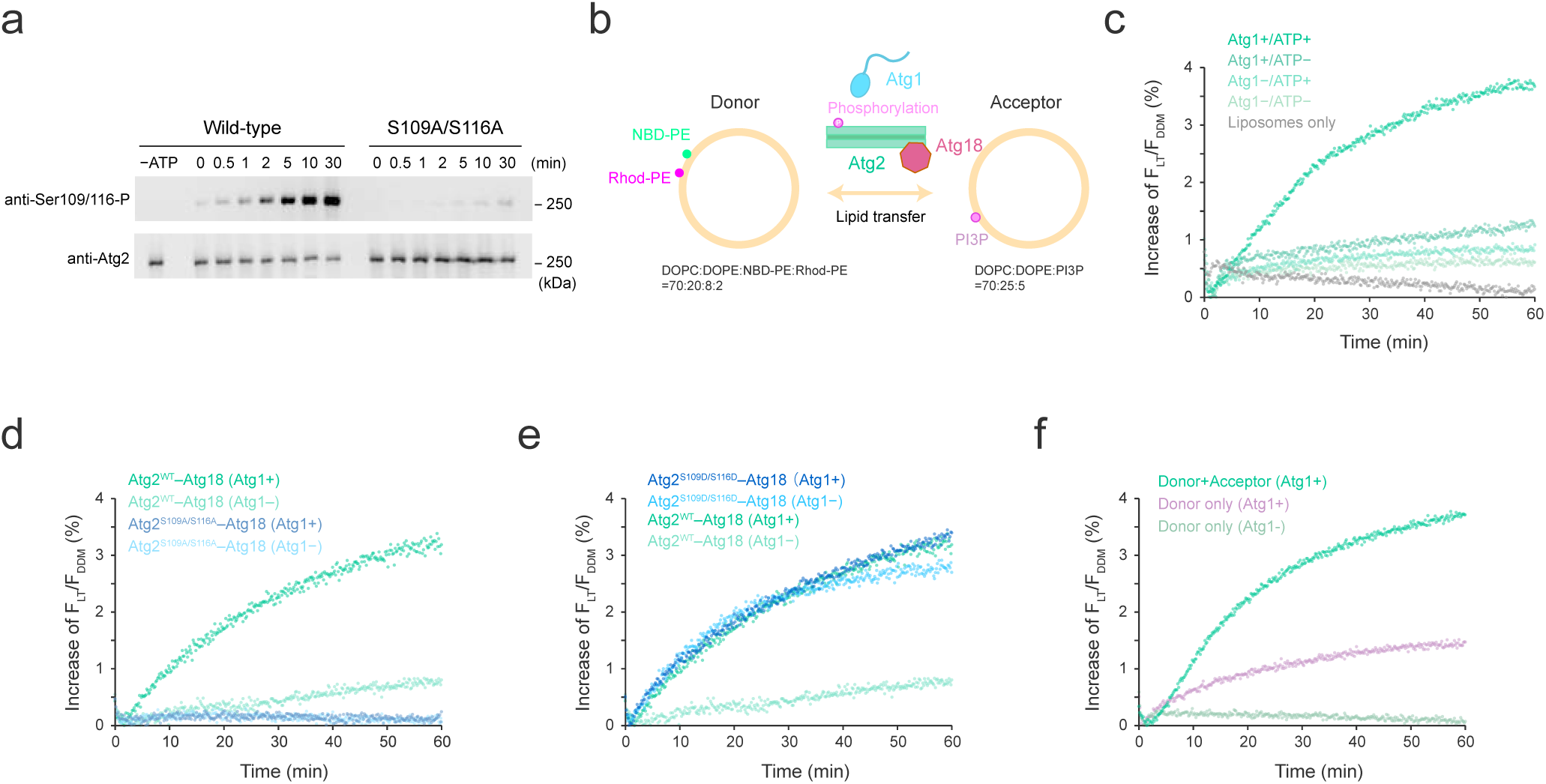
Atg1-mediated phosphorylation enhances the apparent LT signal in a FRET-based assay. **a,** Western blot analysis of Atg1-mediated phosphorylation of Atg2. Phosphorylation of Atg2 was detected using an anti-Ser109/Ser116-P Atg2 antibody. **b,** Schematic representation of the FRET-based LT assay. Donor and acceptor liposomes were mixed with Atg2–Atg18, Atg1, and ATP. Lipid transfer from donor to acceptor liposomes dilutes the FRET pair, NBD-PE and Rhod-PE, resulting in dequenching of NBD fluorescence. **c–f,** FRET-based LT assays. **c,** Atg2–Atg18 in the presence or absence of Atg1 and ATP. **d–e,** Atg2S109A/S116A and Atg2S109D/S116D, respectively, in the presence or absence of Atg1. **f,** Atg2–Atg18 under three conditions: with both donor and acceptor liposomes in the presence of Atg1, or with donor liposomes alone in the presence or absence of Atg1. The increase in NBD fluorescence, F_LT_ was normalized to the maximum fluorescence intensity after addition of DDM, F_DDM_. For all FRET-based LT assays, data are shown as the mean of at least three wells prepared and analyzed in parallel.

However, we noticed that phosphorylation of Atg2 by Atg1 also markedly increased the apparent LT activity in an experimental system containing donor liposomes but no acceptor liposomes (Fig. 4f). This suggests that NBD–Rhod FRET dequenching occurred even in the absence of lipid transfer between donor and acceptor liposomes. Considering the results of the MD simulations, we speculated that phosphorylation of H2 opens the entrance to the Atg2 cavity, allowing NBD-PE and Rhod-PE to enter the cavity and thereby causing dequenching. In this assay, it was not possible to distinguish whether Atg2 transferred lipids between liposomes through the bridge model, merely accommodated lipids within its cavity, altered fluorophore spacing by promoting membrane tethering, or transferred lipids through the shuttle model.

### Narrowing Atg2’s hydrophobic cavity along the conduit impairs autophagy

Next, we decided to examine whether Atg2 actually transports lipids in vivo via the bridge model rather than the shuttle model. Only the bridge model requires lipid movement through the entire cavity, implying that narrowing the cavity along the conduit would disrupt lipid flow (Fig. 5a). Atg2 has five RRs that resemble half-pipes (Fig. 1a), connected by loops across the slit (ARCH1–ARCH4) (Fig. 5b). In addition, two loops (ARCH5, ARCH6) cross the slit around the C-terminal region (CR) (Fig. 5b). We reasoned that deleting or shortening these loops could narrow the cavity without altering the amino acid composition lining the hydrophobic pathway, potentially hindering lipid flow. We generated five Atg2 mutants (ΔARCH1, ΔARCH2+ΔARCH3, ΔARCH4, ΔARCH5, ΔARCH6) in which each loop was deleted or replaced by a short glycine linker (Fig. 5c).

**Figure 5.**
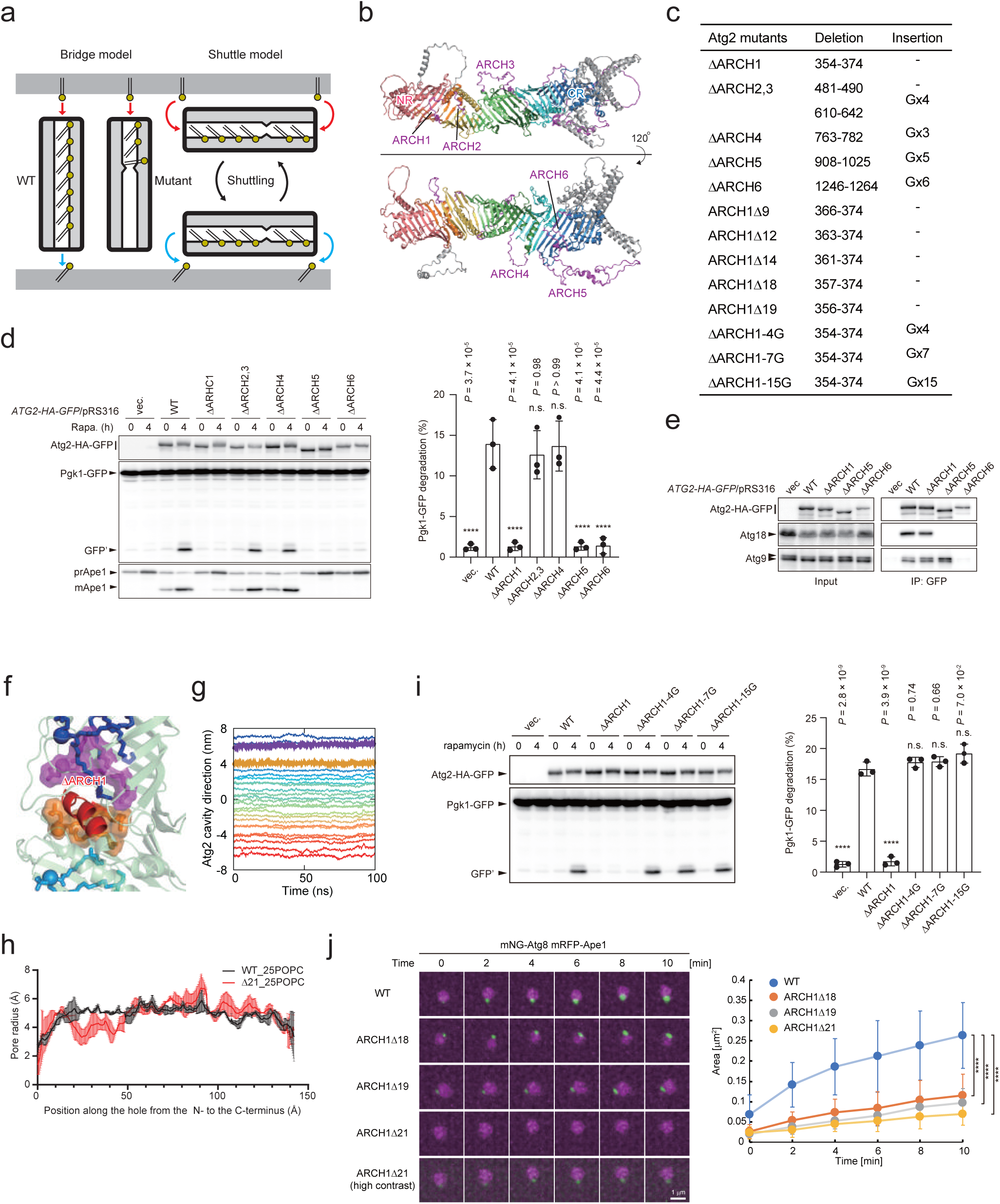
Narrowing the Atg2 cavity along the conduit impairs autophagosome formation. **a,** Schematic drawing of the effect of an internal narrowing of the long cavity on bridge-type and shuttle-type lipid transfer. **b,** Positions of the six ARCHs in the Atg2 structure. **c,** Summary of Atg2 ARCH mutants used in this study. **d,** Effect of ARCH deletion from Atg2 on the autophagic activity assessed by Pgk1–GFP processing and Ape1 maturation. Atg2 levels, Pgk1–GFP degradation, and Ape1 maturation were examined by immunoblotting using antibodies against HA, GFP, and Ape1, respectively. Graph shows the mean ± s.d. (n = 3) of the ratio GFP’/(Pgk1–GFP + GFP’). \*\*\*\**P* < 0.0001; n.s., not significant (Tukey’s multiple-comparisons test; n = 3 independent experiments). **e,** Coimmunoprecipitation analysis of Atg2 with Atg9 and Atg18. **f,** POPC distribution in the narrowed cavity after a 100-ns MD simulation of Atg2^ΔARCH1^ filled with 25 POPC molecules. **g,** Time evolution of the intermolecular distances between POPC molecules along the Atg2 cavity during 100-ns MD simulations of Atg2^ΔARCH1^ filled with 25 POPC molecules. **h,** Pore radius of the cavity for Atg2^WT^ (black) and Atg2^ΔARCH1^ (red) after a 100-ns MD simulation. Error bars indicate s.d. of three independent calculations. **i,** Effect of Gly insertion into the ARCH1-deleted region on the autophagic activity assessed by Pgk1–GFP processing. Graphs show the mean ± s.d. (n = 3) of ratio GFP’/(Pgk1–GFP + GFP’). \*\*\*\**P* < 0.0001; n.s., not significant (Tukey’s multiple-comparisons test; n = 3 independent experiments). **j,** Time-lapse imaging of IMs labeled with mNeonGreen–Atg8 in cells overexpressing Ape1 and treated with rapamycin. The time at which an mNeonGreen–Atg8 punctum first appeared was defined as 0 min, and the size of the IM was measured for up to 10 min. For each strain, we conducted three 1-h imaging sessions at 2-min intervals and quantified 10 IMs in each session. ****P < 1.0 × 10 ¹ for WT versus ARCH1Δ18, WT versus ARCH1Δ19, and WT versus ARCH1Δ21 at 2, 4, 6, 8, and 10 min (Dunn’s multiple-comparisons test).

We measured autophagic activity in *atg2*Δ cells expressing each mutant using Pgk1–GFP cleavage assays. Deleting ARCH2 and ARCH3 together, or deleting ARCH4, had negligible effects, whereas ΔARCH1, ΔARCH5, or ΔARCH6 severely impaired both assays (Fig. 5d). Coimmunoprecipitation revealed that ΔARCH5 lost Atg18 binding and ΔARCH6 lost both Atg18 and Atg9 interactions (Fig. 5e), likely causing their profound autophagy defects. Consistently, AlphaFold2 predicts that segments of ARCH5 and ARCH6 interact directly with Atg18 (Supplementary Fig. 4a). In contrast, ΔARCH1 still bound Atg18 and Atg9 at wild-type levels (Fig. 5e) and localized normally to the PAS (Supplementary Fig. 4b). Notably, only the ΔARCH1 mutant showed a pore radius <4 Å in the AlphaFold models (Supplementary Fig. 4c). Thus, specifically narrowing the cavity by ARCH1 deletion appears to disrupt autophagy.

### A continuous cavity with a radius greater than 4 Å is required for autophagosome formation

We further examined how ARCH1 deletion affects lipid packing in Atg2’s cavity. Even with ΔARCH1, ∼25 phospholipids occupied the cavity after 100 ns MD simulations, but their side-by-side packing was persistently disrupted at the constriction (Fig. 5f, 5g). Quantification showed that part of the pore radius remained <4 Å (Fig. 5h). Based on these observations, we postulated that the autophagic defect arises from an inability of phospholipids to traverse the cavity in single file.

To confirm this, we manipulated how many residues in ARCH1 were removed (9, 12, 14, 18, 19, or all 21). The AlphaFold models predicted that fewer deleted residues would gradually increase the pore radius at ARCH1 from ∼3.7 Å up to ∼5 Å (Supplementary Fig. 4d). Assessing autophagy in cells expressing these mutants (Supplementary Fig. 4e) revealed a direct correlation between the minimum pore radius and autophagic efficiency. Deletions of 18 or 19 residues impaired autophagic activity, albeit less than the full 21-residue deletion, whereas smaller deletions (9, 12, or 14 residues) retained near-wild-type function. Because the residues removed from ARCH1 might themselves have an autophagy-critical function, such as mediating interactions with other proteins, we inserted a glycine repeat into the 21 residue deletion mutant and found that insertion of four or more Gly residues, which expanded the pore radius to >4 Å (Supplementary Fig. 4d), restored autophagic activity to wild-type levels (Fig. 5i). These data strongly indicate that a pore radius >4 Å is essential for normal autophagy, underscoring the importance of an unobstructed Atg2 cavity for side-by-side lipid passage.

We next tested how ARCH1 deletions affect IM expansion. In yeast cells overexpressing Ape1, a selective autophagy substrate that forms large, liquid-like droplets, the IM surrounding these droplets becomes sufficiently large for quantitative fluorescence analysis. In cells expressing Atg2^WT^, we observed the typical cup-shaped IM labeled by mCherry–Atg8 (Supplementary Fig. 4f). By contrast, Atg2^ARCH1Δ^^18^ and Atg2^ARCH1Δ^^19^ displayed a reduced frequency of such IMs, and Atg2^ARCH1Δ^^21^ scarcely showed any cup-shaped structure (Supplementary Fig. 4f). Because this analysis detected end-point structures, we next quantified how the mutations affect the rate of autophagosome formation by time-lapse imaging. Although Atg2’s LT activity is not normally rate-limiting for autophagosome formation, prior work showed that it becomes rate-limiting when another BLTP Vps13 is deleted^5^. Accordingly, in a vps13Δ background we compared wild-type and mutant Atg2 using time-lapse microscopy and found that ARCH1 deletions exhibited a pronounced, statistically significant reduction in the rate of membrane formation (Fig. 5j). These findings confirm that narrowing the hydrophobic cavity restricts lipid flow to the IM and impedes autophagosome biogenesis.

### Narrowing the cavity impairs the flow of lipophilic dye from the ER to the IM in vivo

Visualizing endogenous lipid transport from the ER to the IM in cells has proven technically challenging. Previously, we reported that the fluorescent dye octadecyl rhodamine B (R18), which has a single 18-carbon acyl chain, stains not only the ER but also autophagic structures such as the PAS and the IM^66^. Moreover, *ATG2 deletion* abolished R18 staining of the PAS, indicating that R18 likely travels from the ER to autophagic membranes via Atg2^66^. Recently, our more detailed analyses confirmed that R18 does not migrate by nonspecific free diffusion; rather, its movement from the outer to the inner leaflet of the plasma membrane, through the ER, and ultimately to the IM is driven by the specific actions of a defined set of proteins^67^. Moreover, we confirmed that R18 does not label fully expanded IMs via free diffusion in the cytosol (Supplementary Fig. 5a–5c), and that purified Atg2 binds approximately 15 R18 molecules per Atg2 molecule (Supplementary Fig. 5d). We assessed whether cavity narrowing at ARCH1 impairs this transport. As shown in Fig. 6a, R18 labeled both the ER and the IM in *atg2*Δ cells expressing wild-type Atg2. In contrast, R18 staining at the IM was reduced in cells expressing Atg2^ARCH1Δ^^9^ and barely detectable in those expressing Atg2^ARCH1Δ^^19^ or Atg2^ARCH1Δ^^21^ (Fig. 6a, 6b), even though all mutants showed normal ER staining comparable to wild type. Thus, narrowing the groove impedes R18 (and presumably phospholipid) flow from the ER to the IM, further supporting the physiological relevance of a continuous Atg2 conduit.

**Figure 6.**
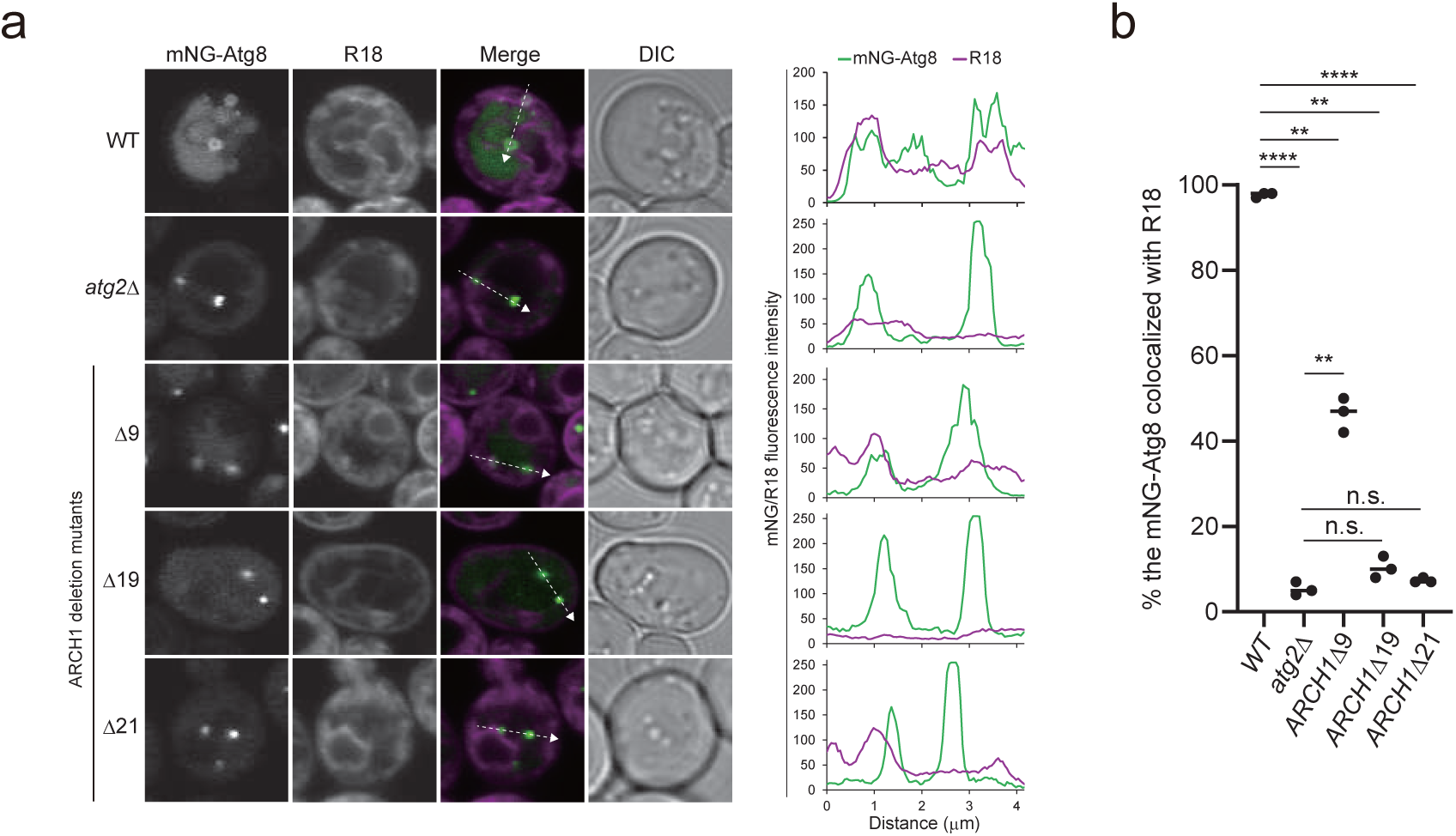
The continuous cavity of Atg2 is important for the ER-to-IM transfer of a lipophilic dye. **a,** Colocalization between mNeonGreen–Atg8 and R18 in cells in the SEY6210 background treated with rapamycin to induce autophagy. Scale bar, 2 μm. Graphs show fluorescence-intensity line profiles from the images on the left. **b,** The percentage of colocalization was calculated from 100 cells per strain across three independent experiments in (**a**); error bars indicate s.d. **P = 0.0052, ***P = 0.00072, ****P = 0.0000069 (WT versus atg2Δ), ***P < 0.00000001 (WT versus atg2Δ21), Welch and Brown–Forsythe test.

### The GUV–SLB system visualizes Atg1-activated bidirectional intermembrane lipid transport through the Atg2 conduit

The in silico and in vivo results suggested that ARCH1 deletion selectively impairs bridge-type lipid transfer by Atg2. We therefore next examined whether this mutation also impairs conventional FRET-based LT activity. We purified Atg2^ΔARCH1^ and verified by negative-stain EM that it retained the same overall rod-like shape as Atg2^WT^ (Supplementary Fig. 6a, 6b). In dynamic light scattering (DLS) experiments, ΔARCH1 did not affect membrane tethering activity, further confirming that the deletion did not affect the structural integrity of Atg2 (Supplementary Fig. 6c). Unexpectedly, the FRET-based LT assay^20,22,23^ revealed no difference between Atg2^WT^–Atg18 and Atg2^ΔARCH1^–Atg18 (Supplementary Fig. 6d–6e). Western blot analysis showed that Atg2^ΔARCH1^ was phosphorylated by Atg1 in a manner similar to that of wild-type protein (Supplementary Fig. 6f), and Atg1-mediated phosphorylation activated the LT activity of Atg2^ΔARCH1^, as it did for Atg2^WT^, regardless of whether Ni-NTA-PE was included in the acceptor liposomes to tether Atg2, which possesses a C-terminal hexahistidine (His)-tag, more tightly to the membrane (Supplementary Fig. 6g–6i). As described above, this assay system could detect lipid transfer between liposomes via the bridge model, lipid transfer via the shuttle model, and incorporation of fluorescent lipids into the Atg2 cavity, but it cannot distinguish among these processes. Thus, it was possible that Atg2 performed non-physiological shuttle-like lipid transfer that does not require a continuous cavity in vitro^15,22,23^, and that this activity was detected for Atg2^ΔARCH1^ in the assay. We therefore decided to establish a new assay system that preferentially detects bridge-model-mediated lipid transfer.

Because Atg2–Atg18 is assumed to bridge the ER and the IM in cells in a defined orientation, interacting with the IM via Atg18-mediated PI3P recognition^13^ and with the ER via Atg2’s N terminus (Figs. 2g and 3c)^14^, we designed a bridge-type LT assay using giant unilamellar vesicles (GUVs) and supported lipid bilayers (SLBs). In this assay, two types of membranes were prepared: GUVs containing NBD-PE and SLBs containing Cy5-PE, PI3P, and Ni-NTA-PE (Fig. 7a). Because Atg2 carried a C-terminal His-tag, Atg2–Atg18 was immobilized on the SLB through both His-tag–Ni-NTA-PE and Atg18–PI3P interactions, thereby orienting Atg2–Atg18 on the SLB. C-terminally His-tagged Atg1 was also immobilized on the same SLB and phosphorylated Atg2 upon addition of ATP. After unbound proteins were washed away to suppress shuttle-type lipid transfer, GUVs were added to the SLB, and fluorescent lipid transfer between the GUVs and SLB was monitored by live imaging.

**Figure 7.**
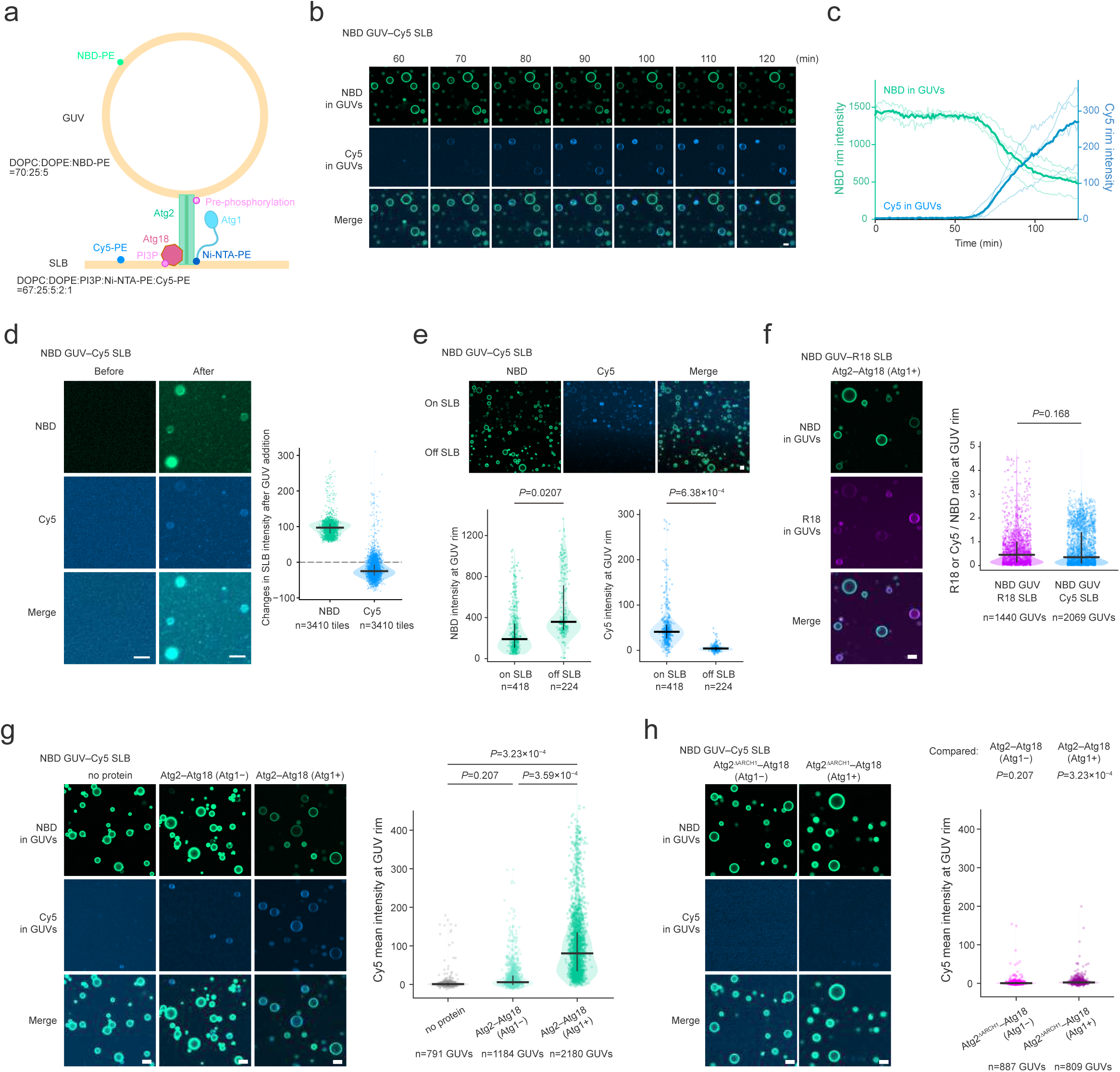
Atg1-mediated phosphorylation activates bridge-type LT activity of Atg2. **a,** Schematic representation of the bridge-type LT assay using GUVs and SLBs. Atg2–Atg18 was immobilized on the SLB through the interaction between the C-terminal His tag of Atg2 and Ni-NTA lipids and phosphorylated by Atg1 on the SLB. Unbound proteins were washed out to suppress shuttle-type lipid transfer. NBD-labeled GUVs were added to Cy5-labeled SLBs to monitor fluorescent lipid transfer between the membranes. **b, c,** Time-lapse imaging of fluorescent lipid transfer between NBD-labeled GUVs and Cy5-labeled SLBs. Representative images are shown in (**b**), and NBD and Cy5 rim intensities of individual GUVs are plotted over time in (**c**). **d,** Changes in NBD and Cy5 fluorescence in the SLB plane before and after GUV addition. Fluorescence changes were quantified using 16 × 16 pixel tiles in GUV-free SLB regions, excluding regions occupied by settled GUVs. **e,** NBD and Cy5 rim intensities of individual GUVs on or off the SLB. **f,** Fluorescent lipid transfer between NBD-labeled GUVs and R18-labeled SLBs. The ratio of R18 or Cy5 rim intensity to NBD rim intensity was quantified for individual GUVs. **g,** Fluorescent lipid transfer between NBD-labeled GUVs and Cy5-labeled SLBs in the absence of proteins, in the presence of Atg2–Atg18 without Atg1, or in the presence of Atg2–Atg18 with Atg1. **h,** Fluorescent lipid transfer between NBD-labeled GUVs and Cy5-labeled SLBs using Atg2^ΔARCH1^–Atg18 in the presence or absence of Atg1. Scale bars, 10 μm in all images. Dots indicate individual GUVs except for (**d**). Horizontal bars indicate the median, and vertical bars indicate the interquartile range. Each field of view was treated as one independent replicate. For each field, the median GUV intensity was calculated. The field-level medians were compared using a two-sided Welch’s unpaired t-test.

Time-lapse imaging revealed that, after approximately one hour of incubation, the NBD-PE fluorescence of some GUVs began to decrease, while Cy5-PE fluorescence from the SLB concomitantly appeared at the rim of the same GUVs (Fig. 7b, 7c). Consistent with this bidirectional fluorescence change at the GUV rim, quantification of the SLB plane showed an increase in NBD fluorescence and a decrease in Cy5 fluorescence after GUV addition (Fig. 7d). In contrast, GUVs that were not in contact with the SLB showed higher NBD-PE fluorescence and markedly lower Cy5-PE fluorescence at the rim than GUVs on the SLB, indicating that the fluorescence changes depended on GUV–SLB contact (Fig. 7e). Similar fluorescence transfer from the SLB to GUVs was observed when R18 was used instead of Cy5-PE (Fig. 7f), showing that the GUV–SLB assay can also detect transfer of R18, a lipophilic dye used in cell-based assays (Fig. 6).

We further characterized the mobility of fluorescent lipids and Atg2 in the GUV–SLB assay by FRAP. Cy5-PE, R18, and Rhod-PE showed fluorescence recovery after photobleaching in the SLB, confirming that these fluorescent lipids were laterally mobile in the SLB (Supplementary Fig. 7a–7d). In contrast, Atg2-SNAP showed little fluorescence recovery on the SLB in either the absence or the presence of Atg1, suggesting that Atg2-SNAP was largely immobile once bound to the SLB (Supplementary Fig. 7c, 7d). Consistently, Atg2-SNAP fluorescence was not detected at GUV rims, indicating that the protein remained stably anchored to the SLB (Supplementary Fig. 7e). The low lateral mobility of Atg2-SNAP may reduce the efficiency with which Atg2 becomes appropriately positioned at GUV–SLB contact sites, potentially accounting in part for the approximately 1-h delay before lipid transfer begins and for the substantial fraction of GUVs in which lipid transfer is not initiated. Whole-GUV photobleaching further showed that SLB-derived fluorescent probes could recover at the GUV rim in R18- or Rhod-labeled SLB assays (Supplementary Fig. 7f–7g), suggesting that fluorescent lipid transfer between the SLB and GUVs is reversible in this assay. However, little fluorescence recovery was observed for both NBD-PE and Cy5-PE when Cy5-labeled SLB was used under the same whole-GUV photobleaching conditions (Supplementary Fig. 7h). The basis for this apparent irreversibility in the presence of Cy5-PE in the SLB remains unclear, but it may reflect disruption of the membrane-bridging configuration of the Atg2 conduit during the assay. Regardless of the underlying mechanism, these results argue against GUV–SLB fusion and formation of a continuous membrane, and instead support fluorescent lipid transfer between two distinct membranes through Atg2.

We next examined the requirement for Atg2–Atg18 and Atg1-mediated phosphorylation. Under conditions in which Atg1-phosphorylated Atg2–Atg18 markedly increased Cy5-PE fluorescence at the GUV rim, Cy5-PE transfer from the SLB to GUVs was minimal in the absence of Atg2–Atg18 or Atg1 (Fig. 7g). When Atg2^ΔARCH1^ was used, Cy5-PE transfer from the SLB to GUVs remained low even after Atg1-mediated phosphorylation (Fig. 7h), indicating that the continuous hydrophobic cavity of Atg2 is important for lipid transfer between GUVs and the SLB. These results suggest that Atg1-mediated phosphorylation activates Atg2 as a lipid-transfer bridge connecting the SLB and GUVs, thereby promoting bidirectional fluorescent lipid transfer between the two membranes. Together with the cellular phenotypes of the Atg2 mutants, these results strongly support the conclusion that Atg2-mediated bridge-type lipid transfer through a continuous lipid-flow conduit is important for autophagosome formation.

## Discussion

Atg2 and Vps13-family BLTPs have been proposed to solve a central problem in organelle biogenesis: how cells move bulk glycerophospholipids fast enough to build or repair membranes at contact sites. Prior structural, biochemical, cellular, and computational studies established the essential foundation for this idea, showing that BLTPs contain elongated hydrophobic cavities or grooves, transfer lipids between membranes, and can, in some contexts, physically bridge organelles^22,23,33,38,39,68–71^. Our work builds on this framework by addressing a stricter mechanistic question: does Atg2-dependent autophagosome biogenesis require lipid passage through an uninterrupted internal route? The Atg2^ΔARCH1^ mutant provides a bridge-specific test because it creates a physical constriction in the middle of the conduit without introducing charges into the hydrophobic pathway. This distinction is important. Hydrophobic-to-charged substitutions have been invaluable for establishing that lipid-binding cavities matter, but they can also perturb lipid extraction, lipid solvation inside the cavity, or shuttle-type carrier activity^39,40^. By contrast, an internal roadblock is expected to selectively penalize mechanisms that require lipids to traverse the full length of Atg2. Thus, the ΔARCH1 series complements earlier charge-substitution studies and substantially strengthens the case for a continuous bridge mechanism.

A second advance is conceptual and methodological. Conventional ensemble FRET-based LT assays were crucial for discovering the LT activity of BLTPs including Atg2, yet our controls underscore that FRET dequenching can reflect processes other than productive intermembrane transfer, including lipid entry into the protein cavity, changes in fluorophore spacing upon tethering, or shuttle-like exchange. This concern is reinforced by a recent VPS13A study, which noted that conventional liposome-based BLTP assays likely permit shuttle-like transfer and emphasized the need for reconstituted systems that distinguish lipid flow through a membrane-bridging conduit from shuttling^36^. The GUV–SLB assay developed here was designed to reduce this ambiguity by orienting Atg2–Atg18 on a PI3P-containing supported membrane, removing unbound protein, and monitoring lipid movement by live fluorescence imaging only after apposed membranes are brought into a defined bridging geometry. In this setting, phosphorylated Atg2 supports bidirectional lipid exchange, whereas Atg2^ΔARCH1^ does not, despite retaining activity in the conventional FRET-based assay. This separation between FRET-detectable activity and GUV–SLB-detectable bridge transport is central to the interpretation of the study: the continuous cavity is not merely important for lipid binding or generic transfer, but for lipid flow across a membrane-bridging Atg2 molecule.

The directionality observed in these assays also provides insight into the intrinsic properties of the conduit. In cells, bidirectional Atg2-dependent R18 exchange between the ER and IM was observed at fully expanded IMs surrounding giant Ape1 condensates^67^. Consistently, the GUV–SLB assay detected bidirectional movement of R18 and other fluorescent lipids through the Atg2 conduit (Fig. 7). Together, these observations indicate that Atg2 itself functions as a fundamentally bidirectional conduit rather than as an intrinsic molecular rectifier. This distinction is important because IM expansion requires net lipid flux to be biased from the ER toward the IM. Thus, although the conduit can support exchange in either direction, additional mechanisms must impose directionality during phagophore growth. Identifying the forces or factors that generate this net ER-to-IM flux remains a major challenge for future studies.

Mechanistically, the data suggest that Atg2 is a signal-gated conduit rather than a constitutively open lipid pipe. A large hydrophobic bridge connecting organelle membranes could be hazardous unless its activity is restricted to the correct site. Recent identification of the Atg2 phospho-FFAT–Scs2 interaction provided one layer of such spatial control by explaining how Atg1 phosphorylation helps anchor Atg2 to the ER at autophagosome formation sites^45,46^. The present study extends this principle by showing that Atg1 also controls the transport competence of the bridge itself: Atg1 phosphorylation of H2 provides an additional layer of spatial control by opening the ER-facing entrance of Atg2, thereby coupling autophagy initiation to donor-side lipid entry into the conduit. The Atg1- and Atg9-dependent phosphorylation of Atg2 in vivo further suggests that activation occurs after Atg2 is recruited to the PAS, where lipid flux is needed for IM expansion. This phosphorylation-dependent gating contrasts with transmembrane-anchored BLTPs, for which all-atom MD simulations showed spontaneous lipid desorption from the donor membrane into the hydrophobic cavity^72^. On the acceptor side, Atg18 does more than target Atg2 to PI3P-enriched autophagic membranes: by promoting local membrane lifting near the C-terminal exit, it may lower the barrier for lipid release into the curved IM rim. This organization is conceptually consistent with the VPS13A–XKR1 complex, in which lipids are delivered directly from the BLTP bridge domain into the acceptor membrane, whereas the nearby scramblase XKR1 equilibrates lipids between leaflets rather than serving as the immediate lipid acceptor^36^. This two-sided regulation provides a simple model in which Atg1 opens the donor gate, Atg18–PI3P engagement licenses acceptor-side release, and the unobstructed Atg2 cavity supports high-capacity lipid movement between the two membranes. Such regulation may bias an intrinsically bidirectional conduit toward productive ER-to-IM transport, although how a sustained net flux is generated remains unresolved. More broadly, our results define a general principle for organelle biogenesis: signaling kinases can activate bulk membrane supply by opening a lipid conduit at a selected contact site.

## Declaration of generative AI and AI-assisted technologies in the writing process

During the preparation of this work, the authors used ChatGPT (OpenAI) to improve the readability and language of the manuscript and OpenAI Codex to assist in the development of custom Python scripts for data analysis. The authors reviewed and edited all AI-assisted text and independently reviewed, tested, and validated all AI-assisted code. The authors take full responsibility for the content of the article and the accuracy of the analyses.

## Acknowledgments

We thank Yuki Ishikawa for assistance with protein preparation and Keisuke Obara for providing a plasmid. This work was supported in part by JSPS KAKENHI Grant Numbers JP23K05715, JP24K01980, JP25K09559 (to Y.Sakai), JP19K16071, JP22K06123 (to T.K.), JP24K09373 (to T.M.), JP24K09358, JP22H05550, JP21K06055 (to K.M.), JP20H05313, JP22H02569 (to K.S.), JP21H05249 (to Y.Sugita), JP24K09392 (to T.N.), JP26K01955, JP23H04923 (to Y.F.), JP20K06532 (to T.O.), JP19H05708, JP23K20044, JP25H01320, JP25H01322 (to H.N.), JP19H05707, JP23K20044, JP23K06667, JP24H00060, JP25H00966, JP25H01320, JP25H01321 (to N.N.N.), CREST, Japan Science and Technology Agency Grant Number JPMJCR20E3 (to K.S., N.N.N.), and grants from the Takeda Science Foundation (to T.O., H.N., N.N.N.). Numerical computations were performed using the Fugaku (hp230252 and hp250019) and HOKUSAI supercomputer systems at RIKEN and the Wisteria/BDEC-01 system at the University of Tokyo.

## Author contributions

Y.Sakai, K.M., T.K., T.O., H.N., and N.N.N. designed the research. K.M., T.O., and N.N.N. performed AlphaFold analysis. Y.Sakai and Y.Sugita performed MD simulations. C.K., J.S., T.K., and H.N. performed budding yeast experiments other than R18. L.H. and K.S. performed budding yeast experiments using R18. T.N. performed MS analysis. T.M., K.M., and Y.F. performed in vitro assays. Y.Sakai, T.K., T.M., K.M., Y.Sugita, T.O., H.N., and N.N.N. analyzed data. Y.Sakai, T.O., and N.N.N. wrote the manuscript with input from all authors. N.N.N. supervised the work.

---

**Supplementary Figure 1. Spiral slit observed in the Atg2 cavity.**

**a,** Surface model (left) and cross section of the hydrophobic cavity (right) of the rod region of Atg2. Coloring is based on the electrostatic potential (blue and red represent positive and negative electrostatic potentials, respectively). ARCHs are shown with green loops. **b,** Comparison of the AlphaFold-predicted and cryo-EM structures of the Atg2–Atg18 complex (PDB: 9T0C). Left, superposition of the overall structures. Right, close-up view of the Atg18 region. **c,** Coimmunoprecipitation experiments using anti-FLAG antibody-conjugated beads and cells expressing Atg18 mutants.

**Supplementary Figure 2. Prediction of membrane interaction by the Atg2–Atg18 complex.**

**a,** Prediction of membrane interaction by the N-terminal region of Atg2 (top) and by Atg18 and the C-terminal portion of Atg2 (bottom) using the PPM server. **b,** Structural comparison of the Atg2–Atg18 interaction before and after MD simulations. **c,** Close-up view of the interaction of the FRRG motif of Atg18 with the IM-like membrane (left, from Fig. 2a and 2c) or with the POPC-only membrane (right, from Fig. 2d). **d,** Positional changes of the 27th and 28th POPC molecules (counted from the N-terminal end) inside the Atg2 cavity during the MD simulation. At 0 ns both lipids were inside the cavity; after 200 ns, the 28th POPC had moved into the C-terminal membrane, and the 27th POPC shifted toward the space vacated by the 28th. **e,** All-atom MD simulation of the Atg2–Atg18 complex bound to a membrane composed of POPC. The spheres indicate the phosphorus atoms of POPCs. **f,** All-atom MD simulation of the Atg2–Atg18 complex bound to the IM-like membrane kept planar by restraints. **g,** Detailed interaction of the N-terminal region of Atg2 with the N-membrane after 200 ns MD simulations in Fig. 2e. POPCs are colored gray for carbon, blue for nitrogen, and red for oxygen. Atg2 is colored green for carbon, blue for nitrogen, and red for oxygen. **h,** Coarse-grained MD simulation of the Atg2–Atg18 complex on vesicles. Top, vesicles with ER and autophagic lipid composition were attached to the N- and C-terminus of the Atg2–Atg18 complex, respectively. Bottom, vesicle with ER lipid composition was attached to the N-terminus of the Atg2–Atg18 complex. In both calculations, 28 POPCs were placed in the Atg2 grooves.

**Supplementary Figure 3. Prediction of the effects of H2 phosphorylation on the N-terminal structure of Atg2 and generation and evaluation of the anti-Ser109/116-P antibody.**

**a,** AlphaFold2 and AlphaFold3 models of the N-terminal region of Atg2 colored according to pLDDT scores. **b,** AlphaFold2 model of the N-terminal region of Atg2^WT^. The side chains of Ser109, Thr114, Ser116, and the adjacent hydrophobic residues are shown in stick representation. **c,** AlphaFold3 predicts that phosphorylation of Ser109, Thr114, and Ser116 induces a large-scale conformational change in H2. **d,** Local curvature of the N- and C-membranes averaged over the 100–200-ns interval of a 200-ns MD simulation. **e,** Detailed view of the interaction between the N-terminal region of phosphorylated Atg2 and the N-membrane. Side chains involved in the interaction with the N-membrane are shown as stick models, and the N-membrane is shown as a surface model. Nitrogen, oxygen, and phosphorus atoms are colored blue, red, and orange, respectively. For lipid molecules, those initially loaded into the cavity are shown with carbon atoms in yellow, whereas those that moved from the N-membrane into the cavity are shown with carbon atoms in gray. **f,** AlphaFold3-predicted models of the Ser109, Thr114, and Ser116 Ala and Asp mutants. **g,** Antibody production and purification strategy. **h, i,** ELISA of affinity-purified antibody before and after nonphosphopeptide depletion against immobilized phosphorylated peptide (**h**) or nonphosphorylated peptide (**i**). Antigens were coated at 5 μg/mL (100 μl/well), and antibody fractions were tested as twofold serial dilutions from 1:1,000 to 1:128,000. Individual values from two technical replicates are shown, and the lines connect their mean values.

**Supplementary Figure 4. Effect of ARCH deletions on the function and pore radius of Atg2.**

**a,** Interaction of ARCH5 and ARCH6 with Atg18 observed in the AlphaFold2 model of the Atg2–Atg18 complex. **b,** Yeast cells expressing Atg2–HA–GFP and Atg17–2×mCherry were treated with rapamycin for 2 h and then observed under a fluorescence microscope. Bar, 5 μm. DIC, differential interference contrast. The graph shows the percentage of Atg17–2×mCherry puncta positive for Atg2–HA–GFP. n.s., not significant (unpaired t-test). **c, d,** Pore radius of the cavity of Atg2 mutants from NR to CR calculated using COOT^49^. **e,** Effect of shaving various residue numbers from ARCH1 on the autophagic activity assessed by Pgk1–GFP processing and Ape1 maturation. Graphs show mean ± s.d. (n = 3) of ratio GFP’/(Pgk1–GFP + GFP’) and ratio mApe1/(mApe1 + prApe1). \**P* < 0.05; \*\*\**P* < 0.001; \*\*\*\**P* < 0.0001; n.s., not significant (Tukey’s multiple comparisons test, n=3 independent experiments). **f,** Observation of IMs labeled with 2xmCherry–Atg8 in cells overexpressing Ape1 and treated with rapamycin for 4 h. Graph shows the quantified size of IMs. The center line of the box represents the median; the bottom and top edges of the box represent the first and third quartiles, respectively; the upper whisker extends to the largest data point within 1.5 times the interquartile range (the height of the box) above the third quartile; the lower whisker extends to the smallest data point within 1.5 times the interquartile range below the first quartile. \*\*\**P* < 0.001; \*\*\*\**P* < 0.0001; n.s. not significant (Dunn’s multiple comparisons test, n=2 independent experiments). n = 271 (WT), 304 (ARCH1Δ21), 262 (ARCH1Δ19), 250 (ARCH1Δ18).

**Supplementary Figure 5. R18 labels the IM through a mechanism other than free diffusion.**

**a,** Schematic illustration of the Ape1 overexpression system, in which the IM expands but fails to close into an autophagosome upon reaching its maximum length. **b, c,** Observation of IMs labeled with mNeonGreen–Atg8 in cells overexpressing Ape1 and treated with rapamycin for 3 h. R18 staining (10 min) was performed prior to rapamycin treatment (**b**) or after 3 h rapamycin treatment (**c**). The percentage indicates the fraction of cells showing R18 colocalization with mNeonGreen–Atg8. It should be noted that in (**c**), the 34 % of instances where R18 and mNeonGreen-Atg8 were colocalized correspond to IMs that had not fully expanded. The scale bar represents 5 μm. **d,** Binding of R18 to Strep-Atg2 was examined using Strep-Tactin XT resin. Strep-Atg13 was used as a negative control.

**Supplementary Figure 6. In vitro activities of purified Atg2 proteins.**

**a,** SDS-PAGE of purified Atg2 WT and ΔARCH1 stained with Coomassie Brilliant Blue. **b,** Electron microscopy images of purified Atg2 stained with 1 mM uranyl acetate were captured using JEM1400 electron microscope (JEOL) equipped with Rio16 camera (Gatan). **c,** Dynamic light scattering of liposomes using Zetasizer Nano S (Malvern) based on the method described previously^22^. The solution containing 400 μM liposomes and 200 nM protein was incubated at room temperature for 8 min before measurement. **d,** FRET-based LT assay for Atg2 WT, ΔARCH21, and ΔARCH18 based on the method described previously^22^. **e,** Simple linear regression on the initial phase of the reaction in (**d**) (at 50 and 100 nM) to study the initial rate of lipid transfer, with the slope proportional to the rate. **f,** Western blot analysis of Atg1-mediated phosphorylation of Atg2^ΔARCH1^. Phosphorylation of Atg2 was detected using an anti-Ser109/Ser116-P Atg2 antibody. **g,** FRET-based LT assay using Atg2^ΔARCH1^–Atg18 in the presence or absence of Atg1. **h,** Schematic representation of the FRET-based LT assay using acceptor liposomes containing Ni-NTA-PE. Ni-NTA-PE was incorporated into acceptor liposomes to tether Atg2–Atg18 to acceptor liposomes through the C-terminal His-tag of Atg2. **i,** FRET-based LT assay using acceptor liposomes with or without Ni-NTA-PE. For all FRET-based LT assays, data are shown as the mean of at least three technical replicates.

**Supplementary Figure 7. FRAP analysis of the bridge-type LT assay using GUVs and SLBs.**

**a,** FRAP analysis of Cy5-labeled SLBs. Cy5-PE exhibited lateral diffusion in the SLB. **b,** FRAP analysis of R18-labeled SLBs. R18 exhibited lateral diffusion in the SLB. **c, d,** FRAP analysis of Atg2-SNAP and Rhod-labeled SLBs in the absence (**c**) or presence (**d**) of Atg1. Rhod-PE showed lateral diffusion in the SLB, whereas Atg2-SNAP showed little fluorescence recovery. **e, f,** Localization and FRAP analysis of Atg2-SNAP and Rhod fluorescence in the GUV–SLB assay. Non-labeled GUVs were added onto Rhod-labeled SLBs coated with Atg2-SNAP–Atg18 pre-phosphorylated by Atg1. Images were acquired at the GUV plane and the SLB plane to examine Atg2-SNAP localization (**e**). Whole GUVs were photobleached, and the recovery of Rhod-PE fluorescence in GUVs was monitored (**f**). **g,** FRAP analysis of NBD and R18 fluorescence in GUVs after whole-GUV photobleaching. NBD-labeled GUVs were added onto R18-labeled SLBs coated with Atg2–Atg18 pre-phosphorylated by Atg1. Whole GUVs were photobleached, and the recovery of NBD-PE and R18 signals in GUVs was monitored. **h,** FRAP analysis of NBD and Cy5 fluorescence in GUVs after whole-GUV photobleaching. NBD-labeled GUVs were added onto Cy5-labeled SLBs coated with Atg2–Atg18 pre-phosphorylated by Atg1. Whole GUVs were photobleached, and the recovery of NBD-PE and Cy5-PE signals in GUVs was monitored. Scale bars, 10 μm in all images. FRAP experiments of SLBs shown in (**a**–**d**) were repeated at least three times with similar results.

**Supplementary Video 1.** All-atom MD simulations of the Atg2–Atg18 complex bound to a planar membrane containing PI3P. This video was used to generate the images shown in Fig. 2a–2c.

**Supplementary Video 2.** All-atom MD simulations of the Atg2–Atg18 complex bound to separate planar membranes. This video was used to generate the images shown in Fig. 2e–2g.

**Supplementary Video 3.** All-atom MD simulations of the H2-phosphorylated Atg2–Atg18 complex bound to separate planar membranes. This video was used to generate the images shown in Fig. 3a–3e.

**Supplementary Data 1.** Mass spectrometry analysis to identify phosphorylation sites of Atg2.

**Supplementary Data 2.** ImageJ macros used for quantitative fluorescence-image analysis.

## Methods

### Structural prediction using AlphaFold2 and AlphaFold3

The Atg2 structure shown in Figs. 1a, 1b, and 5b; Supplementary Figs. 1a and 3a; and the left panel of Supplementary Fig. 3b was downloaded from the AlphaFold Protein Structure Database (model identifier: AF-P53855-F1-v4; https://alphafold.ebi.ac.uk/entry/AF-P53855-F1). The structure of the Atg2–Atg18 complex in Fig. 1d and Supplementary Figs. 1b and 4a was predicted using AlphaFold2 v2.3 installed on a local computer (Sunway Technology Co., Ltd., Tokyo, Japan)^49^. Predictions were run in AlphaFold-Multimer mode^51^, with five models, one seed per model, and default multiple-sequence-alignment generation using the MMseqs2 server^73^. The unrelaxed predicted models were subjected to Amber relaxation, and the relaxed model with the highest confidence based on predicted LDDT scores was selected for MD simulations and figure preparation^49^. The structure of the phosphorylated Atg2–Atg18 complex for MD simulations, the structures of Atg2 (1-614) in Supplementary Figs. 3c and 3f, and the structure of full-length Atg2 in the right panel of Supplementary Fig. 3a were predicted using AlphaFold3 on the AlphaFold Server (https://alphafoldserver.com/)^64^. Structural figures were prepared using PyMOL (http://www.pymol.org/pymol)^74^.

### Plasmid construction

The plasmids used in this study are listed in Supplementary Table 1^75^. Atg1 was constructed in the pFastBac Dual expression vector with an N-terminal SNAP tag and a C-terminal HRV 3C protease recognition site followed by FLAG and His tags. Atg2 and Atg18 mutants were generated by inverse PCR with PrimeSTAR Max DNA Polymerase using pRS316-ATG2–HA–EGFP, pRS426-GAL1pro-ATG2–TS–H8, or pRS315-ATG18–3xFLAG as templates^22^. The integrity of the protein-coding region was confirmed by DNA sequencing.

### Expression and purification

Atg1 was expressed using the Bac-to-Bac baculovirus expression system (Invitrogen) and purified as described previously^76^. Briefly, Sf9 insect cells expressing Atg1 were disrupted by sonication, and the lysates were clarified by centrifugation. Atg1 was purified using HisPur Ni-NTA resin (Thermo Fisher Scientific), followed by size-exclusion chromatography on a HiLoad 26/600 Superdex 200 prep grade column (Cytiva).

Atg2 and Atg18 were expressed in BJ3505 yeast cells lacking endogenous ATG2 and ATG18 genes and purified according to previously published protocols^22^, with minor modifications. Briefly, yeast cells transformed with the pRS426-based plasmids listed in Supplementary Table 1 were disrupted using zirconia beads, and the lysates were clarified by centrifugation. Atg2^WT^, Atg2^S109D/S116D^, and Atg2^S109A/S116A^ were purified using TALON resin (Clontech), followed by Strep-Tactin XT 4Flow high-capacity resin (IBA Lifesciences). Atg2^ΔARCH1^ and Atg2-SNAP were purified using HisPur Ni-NTA resin, followed by Strep-Tactin XT high-capacity resin (IBA Lifesciences). Atg18 was purified using Strep-Tactin XT high-capacity resin, followed by size-exclusion chromatography on a Superdex 200 Increase 10/300 GL column (Cytiva). Purified proteins were concentrated and stored at −80°C until use.

### FRET-based LT assay

1,2-dioleoyl-sn-glycero-3-phosphocholine (DOPC: #850375), 1,2-dioleoyl-sn-glycero-3-phosphoethanolamine (DOPE: #850725), 1,2-dioleoyl-sn-glycero-3-phosphoethanolamine-N-(7-nitro-2,1,3-benzoxadiazol-4-yl) (ammonium salt) (NBD-PE: #810145), 1,2-dioleoyl-sn-glycero-3-phosphoethanolamine-N-(lissamine rhodamine B sulfonyl) (ammonium salt) (Rhod-PE: #810150), 1,2-dioleoyl-sn-glycero-3-phosphoethanolamine-N-(Cyanine 5) (Cy5-PE: #810335), and 1,2-dioleoyl-sn-glycero-3-[(N-(5-amino-1-carboxypentyl)iminodiacetic acid)succinyl] (nickel salt) (Ni-NTA-PE: #790404) were purchased from Avanti Polar Lipids. Phosphatidylinositol 3-phosphate diC16 (PI3P: #P-3016) was from Echelon Biosciences.

The FRET-based LT assay shown in Supplementary Fig. 6d was performed as described previously^22^. The lipid transfer experiment was initiated by adding premixed donor and acceptor liposomes to a cuvette equipped with a stir bar, followed by the addition of the protein. The total volume was 2,000 µL, containing 6–100 nM Atg2, 6–100 nM Atg18, 20 µM donor liposomes, and 100 µM acceptor liposomes. After the measurement was completed, n-dodecyl-β-D-maltoside (DDM) was added to the reaction mixture to a final concentration of 0.4%. After 40 min, the NBD fluorescence intensity in the presence of DDM was measured. A simple linear regression of the initial phase of the reaction was performed to determine the slope (Supplementary Fig. 6e).

The FRET-based LT assay shown in Fig. 4c–4f and Supplementary Fig. 6g, 6i was performed as follows. Donor and acceptor liposomes were prepared separately. Donor liposomes consisted of 70% DOPC, 20% DOPE, 8% NBD-PE, and 2% Rhod-PE. Acceptor liposomes consisted of either 70% DOPC, 25% DOPE, and 5% PI3P or 68% DOPC, 25% DOPE, 5% PI3P, and 2% Ni-NTA-PE. Lipid solutions were prepared in chloroform supplemented with a small aliquot of methanol when necessary to dissolve PI3P. Acceptor liposomes with or without Ni-NTA-PE were used as indicated. Each lipid solution was dried in a separate glass vial under argon gas, and the resulting lipid films were further dried overnight under vacuum. The lipid films were hydrated and vortexed in H buffer (20 mM HEPES-NaOH, pH 7.0, and 150 mM NaCl) supplemented with 1 mM MgCl to obtain heterogeneous liposome suspensions. The liposome suspensions were subjected to three cycles of freeze–thaw and then passed through membrane filters with pore sizes of 400 nm and 100 nm, 21 times each, using a mini-extruder (Avanti Research). After centrifugation at 21,600 × g for 10 min, the supernatant was used as the liposome suspension.

In the FRET-based LT assay, 5 μL of Atg2–Atg18 solution was first added to each well of a 96-well plate (Greiner). Atg2^WT^, Atg2^S109A/S116A^, Atg2^S109D/S116D^, or Atg2^ΔARCH1^ was used as indicated. The reaction was initiated by adding 95 μL of liposome suspension containing donor liposomes, acceptor liposomes, Atg1, and ATP to the well. The final concentrations were 50 nM Atg2–Atg18, 20 μM donor liposomes, 20 μM acceptor liposomes, 100 nM Atg1, and 1 mM ATP. The concentrations of donor and acceptor liposomes were expressed as total lipid concentrations. Fluorescence was measured at approximately 25°C using a Cytation 5 cell-imaging multimode reader (BioTek), with excitation at 460 nm using a 20 nm bandwidth filter and emission at 538 nm using a 20 nm bandwidth filter. The fluorescence intensity of NBD-PE was measured at 10-s intervals for 60 min. After the measurement, dodecyl-β-D-maltoside (DDM) was added to a final concentration of 0.4% to obtain the maximum fluorescence intensity of each sample. The fluorescence intensity was normalized to the maximum fluorescence intensity after DDM addition. The increase in normalized fluorescence intensity was calculated as ΔF_LT_/F_DDM_ = (F_LT_ − F_min_)/F_DDM_ and expressed as a percentage, where F_LT_ is the fluorescence intensity at each time point, F_min_ is the minimum fluorescence intensity during the measurement, and F_DDM_ is the maximum fluorescence intensity after DDM addition.

### Generation and peptide-ELISA validation of the anti-Ser109/116-P antibody

A rabbit polyclonal antibody recognizing Atg2 phosphorylated at Ser109 and Ser116 (anti-Ser109/116-P) was generated by Scrum Inc. (Tokyo, Japan; project 32092). A 16-residue doubly phosphorylated peptide, CFLVK(pS)IQDLTN(pS)MLQ, and the corresponding nonphosphorylated peptide, CFLVKSIQDLTNSMLQ, were synthesized at 91.9% and 86.31% purity, respectively. The phosphopeptide was conjugated to a cationized bovine serum albumin (cBSA) carrier and used to immunize one rabbit intradermally. A preimmune sample was collected before immunization. The rabbit received five intradermal immunizations at 14-day intervals. The final bleed was collected 7 days after the fifth immunization. Twenty ml of antiserum were subjected to affinity purification using the immobilized phosphopeptide. Bound antibodies were eluted with 3 M MgCl_2_, yielding 7 mL of antibody at 0.65 mg/mL (4.55 mg total). This preparation was then processed against the corresponding nonphosphorylated peptide to deplete phosphorylation-independent antibodies. The final antibody preparation was 3 mL at 0.61 mg/mL (1.83 mg total) in PBS containing 0.09% NaN_3_ and was supplied under refrigerated conditions. Peptide specificity was evaluated by ELISA before and after depletion against the nonphosphorylated peptide. The phosphorylated or nonphosphorylated peptide was immobilized at 5 μg/mL in PBS (100 μl/well). Antibody fractions were tested as twofold serial dilutions from 1:1,000 to 1:128,000 in PBS containing 0.05% Tween-20, and wells were blocked with PBS containing 0.2% Tween-20. Absorbance at 412 nm was measured in duplicate. Depletion against the nonphosphorylated peptide preserved reactivity with the phosphopeptide while markedly reducing reactivity with the nonphosphorylated peptide.

### Western blot analysis of Atg1-mediated Atg2 phosphorylation in vitro

A reaction mixture containing 50 nM Atg2–Atg18, 100 nM Atg1, 20 μM donor liposomes, 20 μM acceptor liposomes, and 1 mM ATP in H buffer supplemented with 1 mM MgCl_2_ was incubated at room temperature. Atg2^WT^, Atg2^S109A/S116A^, or Atg2^ΔARCH1^ was used as indicated. Donor liposomes consisted of 70% DOPC, 20% DOPE, 8% NBD-PE, and 2% Rhod-PE, and acceptor liposomes consisted of 70% DOPC, 25% DOPE, and 5% PI3P. Samples were collected at 0, 0.5, 1, 2, 5, 10, and 30 min. The reactions were stopped by mixing the samples with SDS-PAGE sample buffer (Nacalai Tesque), followed by boiling at 95°C. Western blotting was performed using anti-Atg2 and anti-Ser109/116-P Atg2 antibodies. Chemiluminescent signals from the antibodies and fluorescent signals from the molecular-weight marker were detected using an Odyssey M imaging system (LI-COR) and analyzed using Empiria Studio software (LI-COR).

### Bridge-type LT assay using GUVs and SLBs

For the preparation of GUVs, lipid solutions composed of either 70% DOPC, 25% DOPE, and 5% NBD-PE or 75% DOPC and 25% DOPE were prepared in chloroform. Each lipid solution was spread onto a thin sheet of polyvinyl alcohol and dried under vacuum for at least 1 h. The dried lipid film was hydrated in 320 mM sucrose at room temperature for 15 min, and the solution containing GUVs and the sheet was transferred to a plastic tube. The resulting GUV suspension was applied to a syringe filter with a pore size of 5 μm. GUVs retained on the filter were washed with H buffer to remove small vesicles and then recovered as the GUV suspension.

For the preparation of SLBs, SUVs were prepared from lipid films composed of either 67% DOPC, 25% DOPE, 5% PI3P, 2% Ni-NTA-PE, and 1% Cy5-PE; 67% DOPC, 25% DOPE, 5% PI3P, 2% Ni-NTA-PE, and 1% octadecyl rhodamine B (R18: #O246, Invitrogen); or 67% DOPC, 25.9% DOPE, 5% PI3P, 2% Ni-NTA-PE, and 0.1% Rhod-PE. Lipid solutions were transferred to glass vials, dried under argon gas, and further dried overnight under vacuum. The resulting lipid films were hydrated and vortexed in H buffer. The lipid suspensions were subjected to three cycles of freeze–thaw and then sonicated on ice using a hand-held probe sonicator to obtain SUV suspensions. The wells of an 8-well chamber slide (Matsunami) were washed with 50% isopropanol and NaOH. The cleaned wells were then incubated with SUV suspension at 200 μM total lipid concentration for 15 min to form SLBs. The SLBs were passivated with 1 mg/mL BSA for 10 min and washed with H buffer.

Atg2–Atg18 was immobilized on the SLBs through the interaction between the C-terminal His tag of Atg2 and Ni-NTA lipids in the SLBs. The SLBs were incubated with 50 nM Atg2–Atg18, 100 nM Atg1, and 1 mM MgATP for 30 min and washed with H buffer to remove unbound proteins. Atg2^WT^, Atg2^WT^-SNAP, or Atg2^ΔARCH1^ was used as indicated. Atg2-SNAP was labeled with SNAP-Surface Alexa Fluor 488 (New England Biolabs). The GUV suspension was then added to the SLBs. Confocal laser scanning microscopy was performed using a Nikon A1 LFOV confocal microscope with a 20× objective lens at room temperature. Time-lapse images were acquired at 1-min intervals. Snapshot confocal images were acquired approximately 2 h after GUVs were added to the SLB. In FRAP experiments for SLBs, confocal images were acquired every 10 s after photobleaching with 100% laser power for 2 s. In FRAP experiments for GUVs, confocal images were acquired every 5 s after photobleaching with 100% laser power for 2 s. Confocal images were acquired using NIS-Elements software (Nikon) and processed using Fiji software. Fluorescence intensities of individual GUVs and SLBs were quantified after background subtraction. The resulting data were analyzed using custom Python scripts developed with assistance from OpenAI Codex (OpenAI). All AI-assisted code was reviewed and validated by the authors.

### Atg2–R18 binding assay

Binding of R18 to Atg2 was examined using Strep-Tactin XT resin. Ten microliters of Strep-Tactin XT 4Flow high-capacity resin was equilibrated with binding/wash buffer containing 20 mM HEPES-KOH, pH 7.5, 500 mM NaCl, and 10% glycerol. The resin was blocked with 1% BSA for 10 min at room temperature to reduce nonspecific adsorption, and then washed extensively with binding/wash buffer. Strep-tagged Atg2 was immobilized on the resin by incubating the resin with 0.5 µM purified Strep-Atg2 for 15 min at room temperature with gentle rotation. Resin-only and Strep-Atg13-bound resin were used as a background and a negative control, respectively. Strep-Atg13 was prepared as described previously^10^. After removal of the unbound fraction, the resin was washed three times with binding/wash buffer. R18 was diluted to 20 µM in binding/wash buffer containing 0.02% Tween-20 and added to the resin. Samples were incubated for 10 min at room temperature in the dark with gentle rotation. The resin was then washed five times with binding/wash buffer to remove unbound R18. Bound material was eluted by incubating the resin with elution buffer, consisting of binding/wash buffer supplemented with 5 mM desthiobiotin, for 10 min at room temperature. R18 in the eluate was quantified by fluorescence measurement using a microplate reader with excitation at 560 ± 10 nm and emission at 590 ± 10 nm. Background fluorescence from resin-only or Strep-Atg13 control samples was subtracted. The amount of Atg2 in the eluate was estimated by SDS-PAGE followed by total protein staining, using purified Atg2 standards loaded on the same gel. The molar ratio of R18 to Atg2 was calculated from the background-subtracted R18 concentration and the Atg2 concentration in the eluate.

### Pgk1–GFP cleavage assay

Yeast strains used in this study are summarized in Supplementary Table 2^77–79^. Samples for analyzing autophagic degradation of Pgk1–GFP and Ape1 maturation were prepared as described previously^14^. For immunoblotting, antibodies against GFP (11814460001; Roche), FLAG (M2, F1804; Sigma-Aldrich), Ape1 (anti-API-2)^80^, and HA (3F10; Roche) were used.

### Immunoprecipitation

In Figs. 1h and 5e and Supplementary Fig. 1c, yeast cells were disrupted in IP buffer [20 mM HEPES-KOH (pH 7.2), 150 mM NaCl, 10% glycerol] containing a 2× Complete protease inhibitor cocktail (Roche) using a Multi-beads Shocker (Yasui Kikai) with 0.5-mm YZB zirconia beads. The lysates were then treated with 0.1% CHAPS while rotating at 4°C for 30 min. The lysates were cleared by successive centrifugations at 15,000 × g for 10 min and 30 min. The supernatants were incubated with GFP-nanobody or anti-FLAG antibody-conjugated beads at 4°C for 90 min. The beads were washed three times with IP buffer containing 0.1% CHAPS. Proteins bound to GFP-nanobody-conjugated beads were eluted by boiling the beads in elution buffer 1 prepared by diluting 5× sample buffer [250 mM Tris-HCl (pH 7.5), 10% SDS, 40% glycerol] 5-fold with IP buffer containing 0.1% CHAPS and 20 mM DTT for 3 min. Proteins bound to anti-FLAG antibody-conjugated beads were eluted by incubating the beads in elution buffer 1 at 65°C for 10 min, and 20 mM DTT was added to the eluates after removing the beads. In Fig. 3h, 3i, yeast cells were disrupted in IP buffer containing a 2× Complete protease inhibitor cocktail, PhosSTOP (Roche), and 500 nM microcystin LR and 50 mM NaF using a Multi-beads Shocker with 0.5-mm YZB zirconia beads. The lysates were then solubilized with 0.2% Triton X-100 while rotating at 4°C for 30 min and cleared as described above. The supernatants were mixed with anti-FLAG antibody-conjugated beads at 4°C for 90 min. After three washes with IP buffer containing 0.2% Triton X-100, bound proteins were eluted by incubating at 65°C for 10 min in elution buffer 2, which was prepared by diluting 5× sample buffer with IP buffer containing 0.2% Triton X-100. After removal of the beads, 20 mM DTT was added to the eluates. The samples were subjected to immunoblotting analysis using antibodies against HA (3F10; Roche), FLAG (M2; Sigma), Atg2 (anti-Apg2)^81^, phospho-Atg2 (anti-Ser109/116-P), Atg9 (anti-Atg9 N-pep)^82^, and Atg18 (anti-18-2)^13^.

For mass spectrometry analysis, yeast cells treated for 4 h with β-estradiol for induction of constitutively active Atg1 expression and rapamycin were disrupted in IP buffer containing a 2× Complete protease inhibitor cocktail, PhosSTOP, and 500 nM microcystin LR and 50 mM NaF using a Multi-beads Shocker with 0.5-mm YZB zirconia beads. The lysates were then solubilized with 0.2% Triton X-100 at 4°C for 30 min and cleared as described above. The supernatants were incubated with anti-FLAG M2 Affinity Gel (Sigma) at 4°C for 2 h. After being washed three times with IP buffer containing 0.2% Triton X-100, bound proteins were eluted by boiling the beads in elution buffer 3 [50 mM HEPES-NaOH (pH 8.0), 1% SDS] for 3 min.

### Mass spectrometry

Sample preparation for LC-MS analysis was performed based on the SP3 method^83^. Eluate samples in elution buffer 3 were reduced with 10 mM DTT for 30 min at 37°C, and subsequently alkylated with 20 mM iodoacetamide for 30 min at room temperature in the dark. Sera-Mag SpeedBead Carboxylate-Modified [E3] Magnetic Particles (Cytiva) and an equal volume of ethanol to the sample were added to the samples, followed by mixing at 1,000 rpm for 5 min. The beads were washed three times with 80% ethanol, resuspended in 50 mM ammonium bicarbonate containing 4 ng/µl Asp-N (Promega) alone or in combination with 8 ng/µl Trypsin/Lys-C Mix (Promega), and incubated overnight at 37°C. After removal of the beads, the supernatants were acidified by the addition of 0.5% trifluoroacetic acid (TFA). Samples were loaded onto GL-Tip SDB cartridges (GL Sciences), washed twice with binding buffer (0.1% TFA, 2% acetonitrile), and eluted with elution buffer 4 (0.1% TFA, 80% acetonitrile). Eluted peptides were dried using the centrifugal concentrator, redissolved in binding buffer, clarified by centrifugation at 20,000 × g for 5 min at 4, and transferred to an LC-MS vial.

LC-MS/MS measurements were performed using an Easy-nLC1000 nanoflow liquid chromatography system and a Q-Exactive tandem mass spectrometer (Thermo Fisher Scientific). The trap column used for nano-HPLC was a 2 cm × 75 μm capillary column packed with 3 μm C18 silica particles (Thermo Fisher Scientific, Cat No. 164946), and the separation column was a 12.5 cm × 75 μm capillary column packed with 3 μm C18 silica particles (Nikkyo Technos). The nano-HPLC flow rate was set to 300 nL/min. Separation was conducted under a 10% to 40% linear acetonitrile gradient over 30 minutes in the presence of 0.1% formic acid. Data were acquired in data-dependent acquisition (DDA) mode controlled by Xcalibur 4.0 (Thermo Fisher Scientific). The DDA settings were as follows: resolution of 70,000 for full MS scans and 17,500 for MS2 scans; AGC target of 3.0E6 for full MS scans and 5.0E5 for MS2 scans; maximum ion injection time (IT) of 60 msec for both full MS and MS2 scans; full MS scan range of m/z 310–1,500; and selection of the top 10 signals in each full MS scan were selected for MS2 scans. DDA measurements were performed twice in each sample.

Data analysis was performed using FragPipe platform equipped with the MSFragger search engine (utilizing the LFQ-Phospho workflow)^84^. The amino acid sequence database for the peptide search was obtained from the UniProt database (UP000002311, downloaded on 27 March 2024).

### Fluorescence microscopy of yeast cells

Fluorescence microscopy for Fig. 5j and Supplementary Fig. 4b, 4f was performed using a DeltaVision Elite microscope system (GE Healthcare, Chicago, IL), as described previously^85^. In Fig. 5j, yeast cells were immobilized on glass-bottom dishes coated with Concanavalin A. Following treatment with rapamycin, time-lapse imaging was performed at 2-min intervals for 1 h. At each time point, 25 z-stack images were acquired, followed by deconvolution and maximum-intensity projection using SoftWoRx 7.0.0 (GE Healthcare). Cells in which fluorescent puncta appeared during the imaging period were selected for further analysis. The area of the isolation membrane at each time point was quantified using ImageJ Script S1 (Supplementary Data 2). The time point at which the fluorescent puncta first appeared was defined as 0 min, and changes in isolation membrane area were analyzed during the following 10 min. Twenty-five (Supplementary Fig. 4f) or ten (Supplementary Fig. 4b) z-stack images at 0.2-μm intervals were captured and deconvolved using SoftWoRx 7.0.0. A maximum-intensity projection was generated using Fiji. The perimeter of the IM was measured using ImageJ Script S2 (Supplementary Data 2). The length of the IM was taken as half of its perimeter. PAS localization was examined using ImageJ Script S3 (Supplementary Data 2).

Fluorescence microscopy of cells treated with R18 was performed as follows. Cells were cultured in SDCA medium to a density of about 5 × 10 cells/mL, and stained with 5 μg/mL of R18 (1 mg/mL stock dissolved in dimethyl sulfoxide) for 10 min at 30°C. After the cells were washed three times with fresh medium, autophagy was induced by the addition of rapamycin (0.2 μg/mL). Cells were harvested, spun at RT with a microcentrifuge, and imaged using an FV-3000 confocal laser-scanning microscope (Olympus) equipped with a UPLSAPO60XO oil-immersion objective (NA 1.42; Olympus). The 488-nm and 561-nm lasers were used for sequential excitation, and fluorescence was recorded at 500–540 nm for mNeonGreen and 560–600 nm for octadecyl rhodamine B (R18; Invitrogen), respectively, using a resonant scanner. Samples were prepared using 76 × 26 mm glass slides (S1225, Matsunami) and 18 × 18 mm coverslips (No. 1-S, Matsunami). Images were analyzed by ImageJ (https://fiji.sc/).

### Molecular dynamics simulations

The initial structure of the Atg2–Atg18 complex was predicted using AlphaFold2. For simulations of PC lipids in the Atg2 cavity, an Atg2 construct lacking the C-terminal segment was used. The initial states of POPC lipids in the Atg2 cavity were first equilibrated by coarse-grained (CG) molecular dynamics simulations based on the MARTINI model (version 2.2)^86,87^. The equilibrated structures were then converted to all-atom models and solvated in a 28 nm × 28 nm × 28 nm box containing TIP3P water molecules and 0.15 M KCl ions. For simulations of the Atg2–Atg18 complex on membranes, the full-length structure of the Atg2–Atg18 complex and membranes containing 900 lipid molecules per leaflet were used. The composition of the autophagic membrane was modeled on the lipid composition of Atg9 vesicles^9^, and was set to POPC:POPE:POPS:POPI:POPI13:POPA = 370:54:63:318:80:15. The location of the Atg2–Atg18 complex on the membranes was predicted using the PPM server^88^. The composition of the ER membrane was set to POPC:POPE:POPS:POPI:POPA:POCL1 = 450:216:81:108:36:9^89^. Twenty-eight POPC molecules were placed in the Atg2 grooves and the initial states of the POPC molecules were prepared by converting the coarse-grained equilibrium states to all-atom models, as described above. The protein and lipids were solvated in a 25 nm × 25 nm × 30 nm box with TIP3P water molecules and 0.15 M KCl ions.

The coarse-grained simulations were performed using GROMACS (version 2022)^90^ and the MARTINI model (version 2.2)^86,87^. In the simulation of lipids in the Atg2 cavity, the initial states of POPC molecules were arranged at equal intervals and in the same orientation in the Atg2 cavity. The protein and lipids were solvated in a 28 nm × 28 nm × 28 nm box with coarse-grained waters and 0.15 M NaCl ions. In the simulations of the Atg2–Atg18 complex on vesicles, 30-nm-diameter vesicles with autophagic and ER lipid compositions were attached to the C- and N terminus of the Atg2–Atg18 complex, respectively. Twenty-eight POPC molecules were placed in the Atg2 grooves, and the tertiary structure of the proteins was maintained using an elastic network model. The proteins and lipids were solvated in a 90 nm × 90 nm × 90 nm box with coarse-grained waters and 0.15 M NaCl ions. The initial configurations were built using the Martini Maker module of the CHARMM-GUI server^91^. Each system was first energy-minimized and equilibrated using the Berendsen thermostat and barostat^92^ followed by 1-μs production runs with a 20-fs time step along with positional restraints on the protein backbone beads with a force constant of 20 kJ mol^-1^ nm^-2^. System temperature and pressure during the production phase were maintained at 303.15 K and 1 atm with the velocity-rescaling thermostat and the semi-isotropic Parrinello–Rahman barostat^93^, respectively. The coarse-grained equilibrium states were converted to all-atom models using the Martini All-atom Converter module of the CHARMM-GUI server^91^.

The all-atom simulations were performed using GENESIS^94^ and the CHARMM36 force field^95^. For the phosphorylated system, the AlphaFold3 model of the Atg2–Atg18 complex was used, with Atg2 Ser109 and Ser116 parameterized as SP2 and Thr114 as THPB. The initial configurations were built using the Membrane Builder module of the CHARMM-GUI server^91^. Energy minimization was performed for 1,000 steps by the steepest-descent algorithm and then by the conjugate-gradient algorithm. A 250-ps NVT simulation was performed at 303.15 K for solvent equilibration, followed by a 1.6-ns NPT equilibration to 1 atm using the Langevin thermostat/barostat^96^. The production MD simulations were performed with a 2-fs time step and Langevin thermostat/barostat. The cavity-loading and Atg2^ΔARCH1^ systems were simulated for 100 ns; the 25-POPC wild-type system was extended to 200 ns for the intermolecular-distance and MSD analyses, and all Atg2–Atg18 membrane systems were simulated for 200 ns. Long-range electrostatic interactions were treated by the particle-mesh Ewald method^97,98^. The short-range electrostatic and van der Waals interactions both used a 12-Å cutoff. All bonds were constrained by the SHAKE/RATTLE algorithm^99,100^. In the planar-membrane control, the phosphorus atoms of the membrane lipids were positionally restrained along the membrane normal with a force constant of 0.1 kcal mol^-1^ Å^-2^.

Simulation trajectories were analyzed using GENESIS. The lipid positions and mean-square displacements (MSDs) in the Atg2 cavity were calculated after translational and rotational fitting to the initial structure to remove overall protein motion during the trajectory. The system was considered equilibrated after 100 ns based on the lipid positions shown in Fig. 1m. The diffusion coefficient was obtained by fitting the MSD over 10–100 ns using the one-dimensional Einstein relation, MSD(t) = 2Dt. The average height of the membrane around the protein was determined by fitting the position of the phosphorus atoms of the lipids at 10-ns intervals from 100 to 200 ns using a fifth-order polynomial function, and the curvature was calculated from the fitted function. Three independent calculations were performed for each simulation to confirm reproducibility.

