## Supplementary Figures 1-7, Supplementary Tables 1,2 for "Atg1 phosphorylates Atg2 to gate phospholipid flow through a continuous conduit for autophagosome biogenesis"

### Supplementary Figure 1

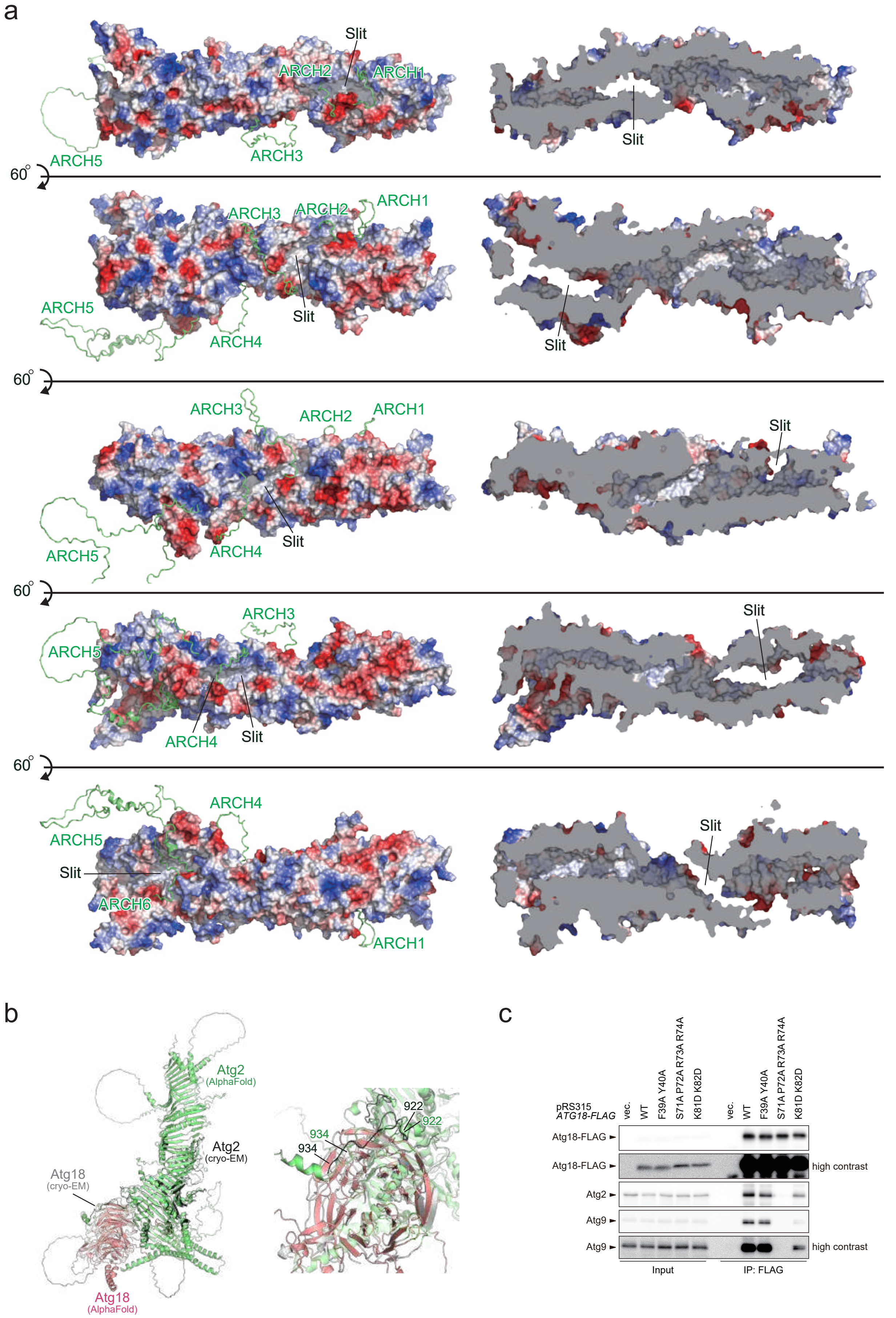

### Supplementary Figure 2

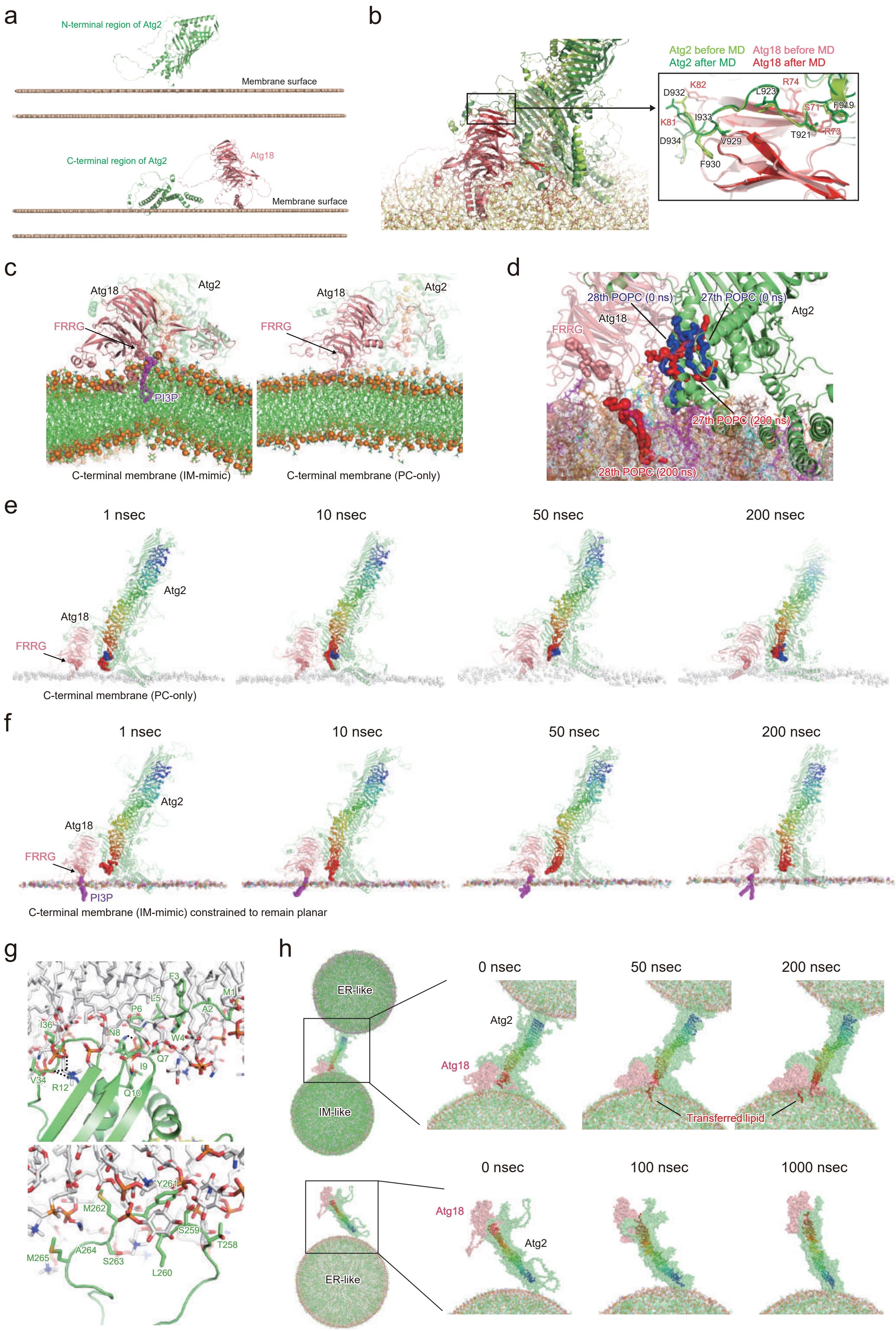

### Supplementary Figure 3

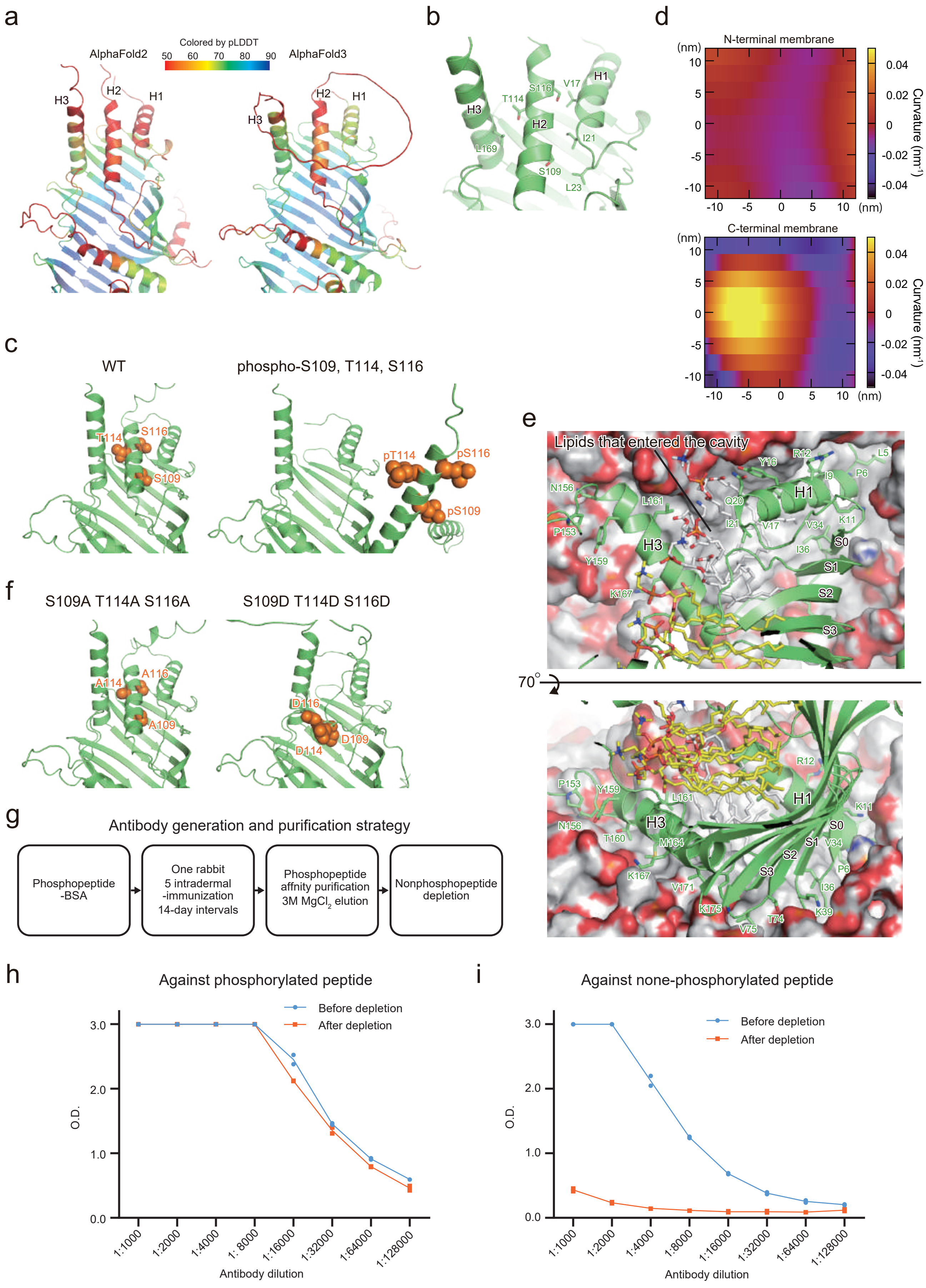

### Supplementary Figure 4

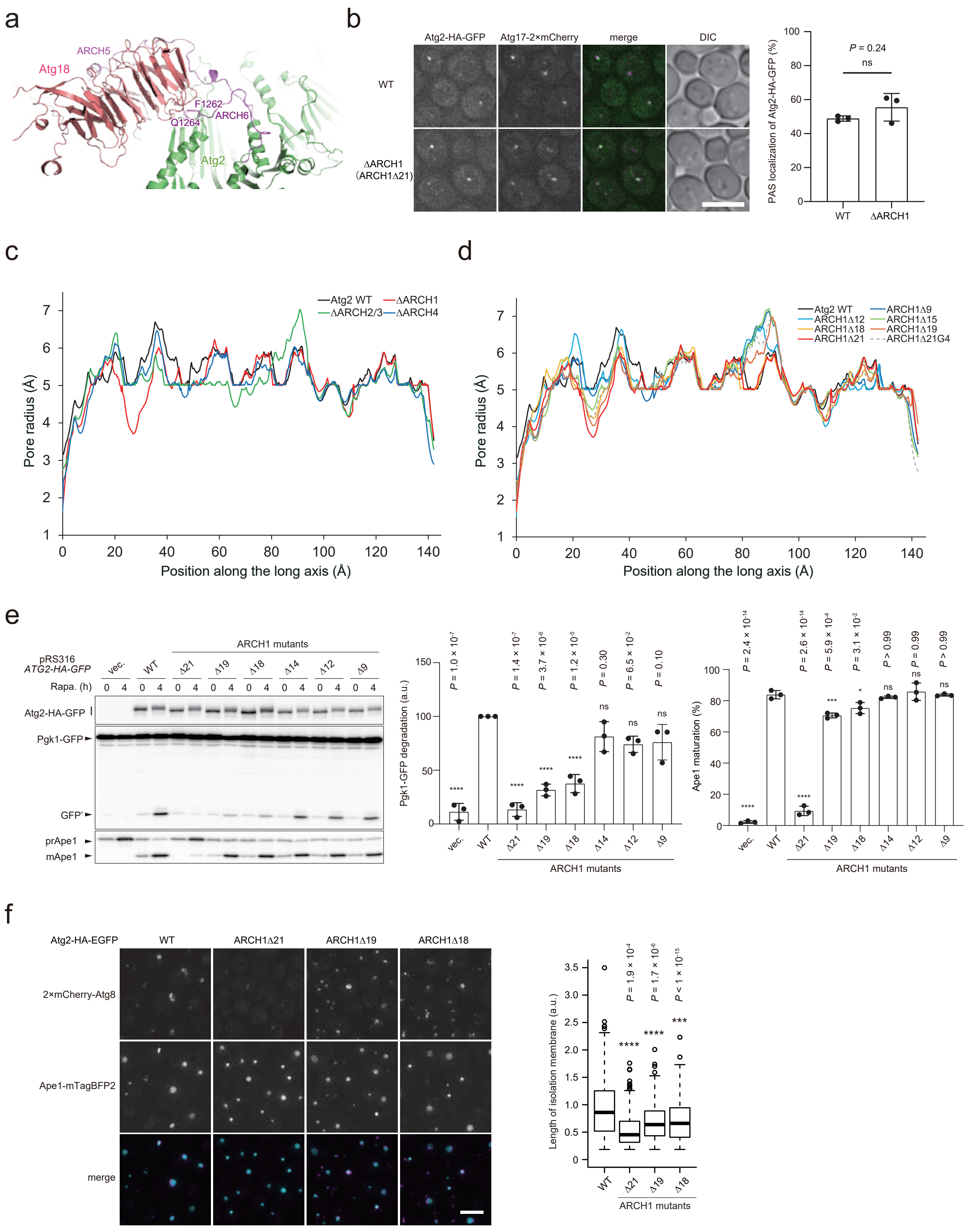

Supplementary Figure 5

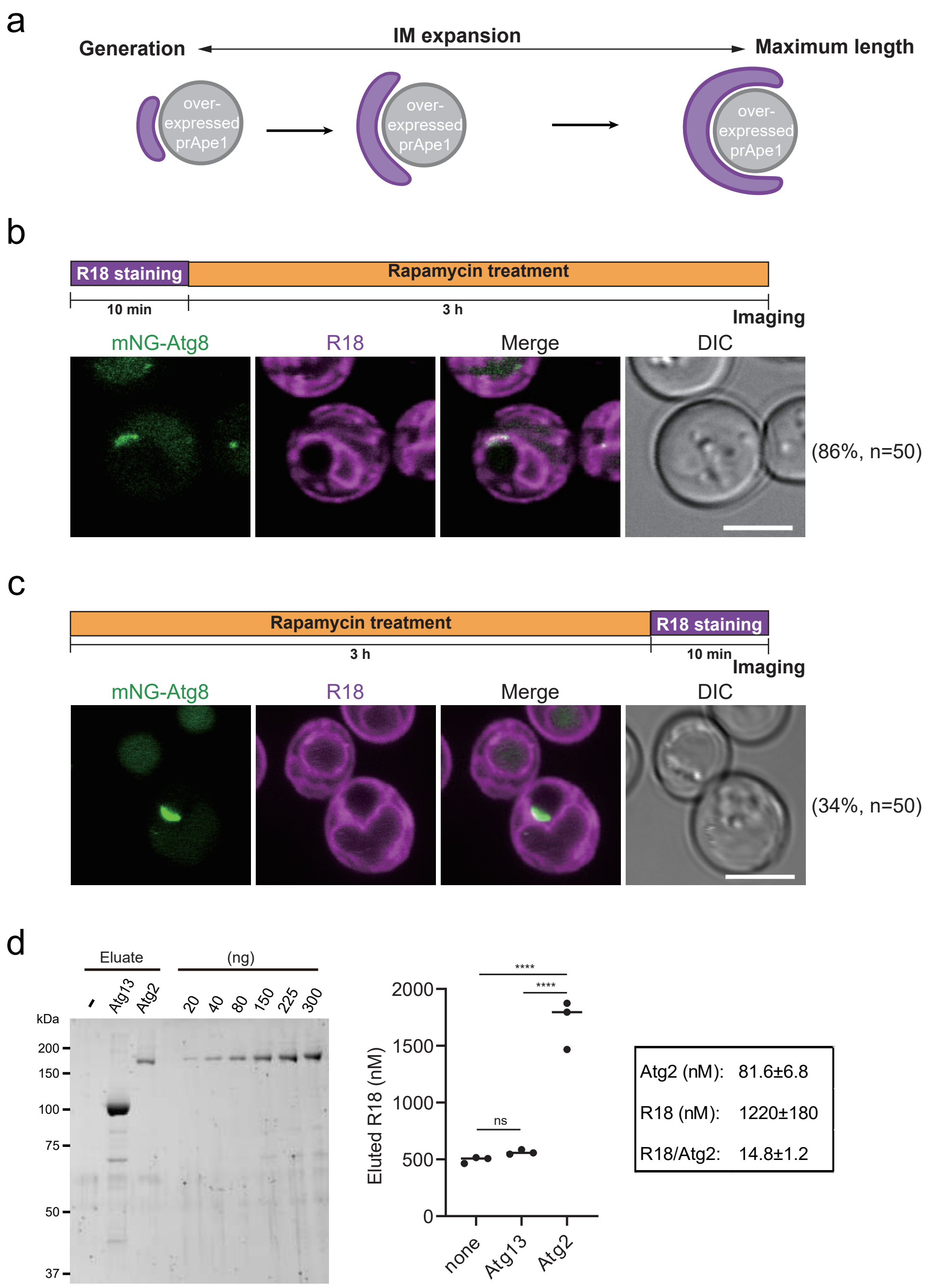

### Supplementary Figure 6

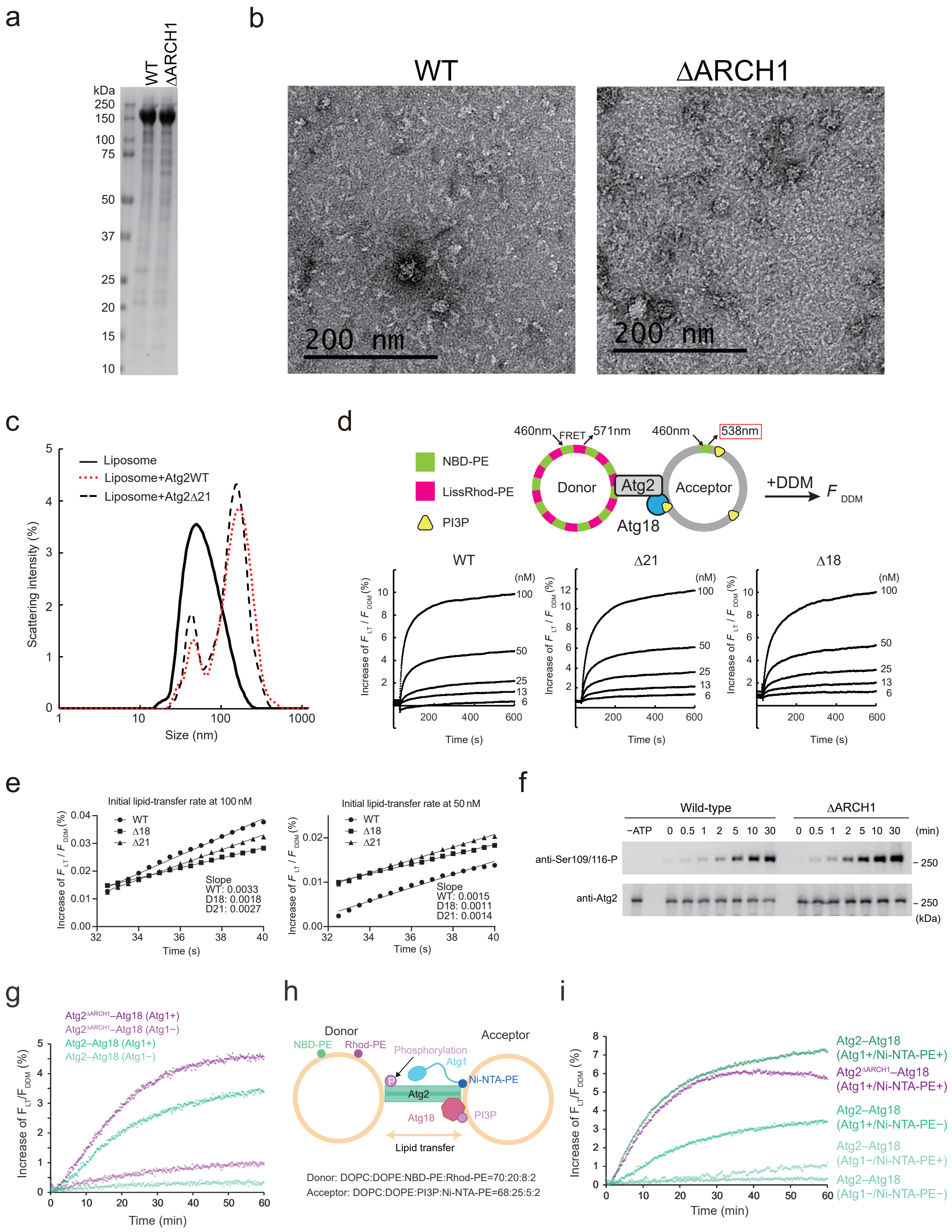

### Supplementary Figure 7

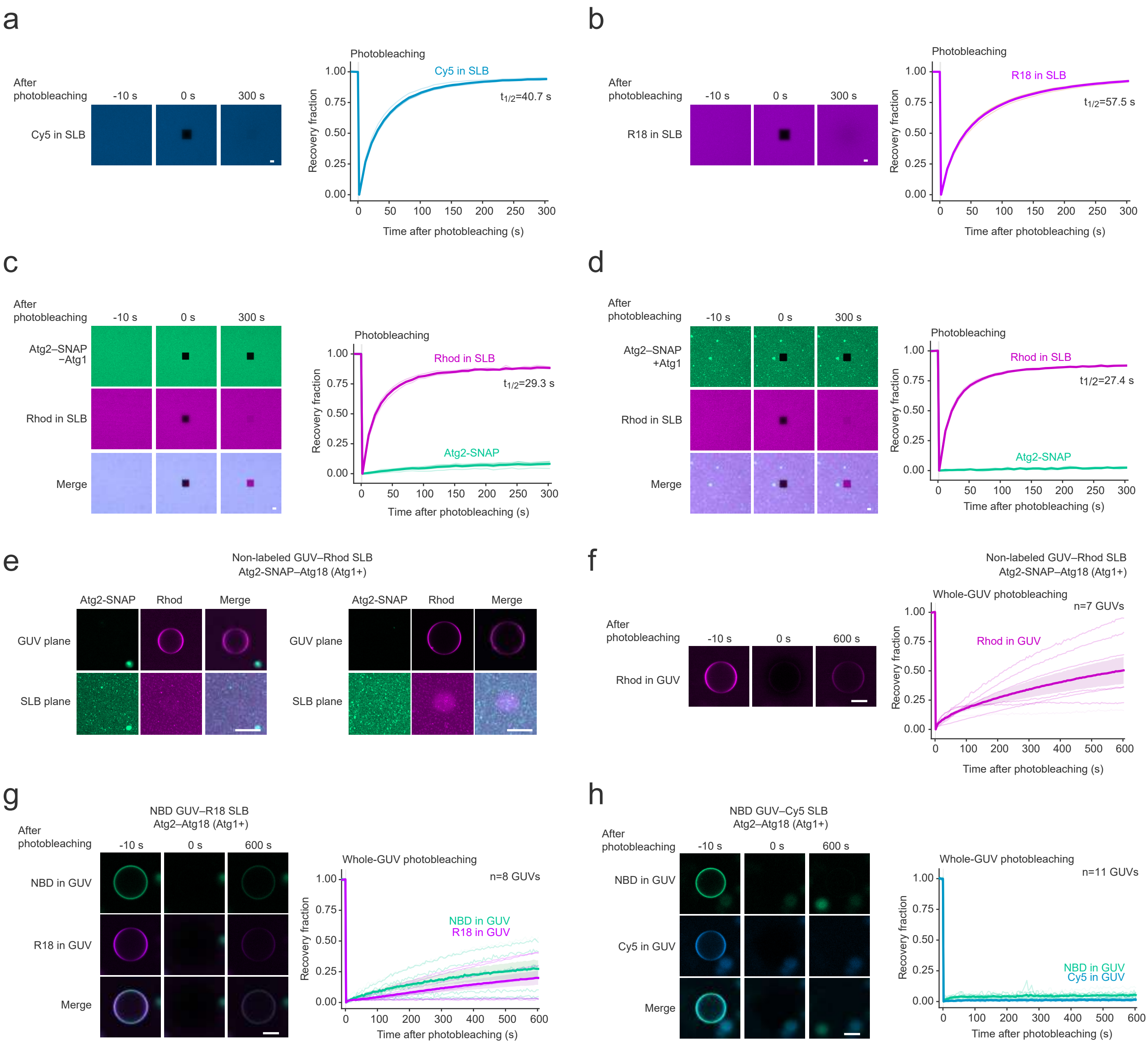

Supplementary Table 1. Plasmids used for this study.

| ID | Plasmid | Source | Mutant name |
|---|---|---|---|
| | pRS316-ATG2-HA-EGFP | Ref 13 | |
| | pRS426-GAL1pro-ATG2-TS-H8 | Ref 13 | |
| pOK101 | pRS315-ATG18 | This study | |
| p2KM-101 | pRS316-ATG2(delta-S354-S374)-HA-EGFP | This study | ΔARCH1/ARCH1Δ21 |
| p2KM-104 | pRS316-ATG2 (S762-GGG-P782)-HA-EGFP | This study | ΔARCH4 |
| p2KM-105 | pRS316-ATG2(L907-GGGGG-D1026)-HA-EGFP | This study | ΔARCH5 |
| p2KM-106 | pRS316-ATG2(G1245-GGGGGG-K1265)-HA-EGFP | This study | ΔARCH6 |
| p2KM-116 | pRS316-ATG2( M480-T491, K610-GGGG-S643)-HA-EGFP | This study | ΔARCH2,3 |
| p2KM-122 | pRS316-ATG2(delta-S356-S374)-HA-EGFP | This study | ARCH1Δ19 |
| p2KM-123 | pRS316-ATG2(delta-C357-S374)-HA-EGFP | This study | ARCH1Δ18 |
| p2KM-124 | pRS316-ATG2(delta-P361-S374)-HA-EGFP | This study | ARCH1Δ14 |
| p2KM-125 | pRS316-ATG2(delta-Q363-S374)-HA-EGFP | This study | ARCH1Δ12 |
| p2KM-126 | pRS316-ATG2(delta-D366-S374)-HA-EGFP | This study | ARCH1Δ9 |
| p2KM-128 | pRS426-GAL1pro-ATG2(delta-S354-S374)-TS-H8 | This study | ΔARCH1/ARCH1Δ21 |
| p2KM-136 | pRS426-GAL1pro-ATG2(delta-C357-S374)-TS-H8 | This study | ARCH1Δ18 |
| p2KM-137 | pRS316-ATG2(F919A T921A L923A)-HA-EGFP | This study | |
| p2KM-138 | pRS316-ATG2(V929A F930A I933A)-HA-EGFP | This study | |
| p2KM-139 | pRS316-ATG2(D932K D934K)-HA-EGFP | This study | |
| p2KM-140 | pRS316-ATG2(E353-GGGG-S375)-HA-EGFP | This study | ΔARCH1-4G |
| p2KM-141 | pRS316-ATG2(E353-GGGGGGG-S375)-HA-EGFP | This study | ΔARCH1-7G |
| p2KM-142 | pRS316-ATG2(E353-Gx15-S375)-HA-EGFP | This study | ΔARCH1-15G |
| p18KM1 | pRS315-ATG18-3xFLAG | This study | |
| p18KM2 | pRS315-ATG18(F39A Y40A)-3xFLAG | This study | |
| p18KM3 | pRS315-ATG18(S71A P72A R73A R74A)-3xFLAG | This study | |
| p18KM4 | pRS315-ATG18(K81D K82D)-3xFLAG | This study | |
| | pYEX-BX[prApe1] | Ref 10 | |
| pYYY59 | pRS316-ATG2-FLAG | Ref 73 | |
| pTKO756 | pRS316-ATG2(S109A)-FLAG | This study | |
| pTKO757 | pRS316-ATG2(S116A)-FLAG | This study | |
| pTKO786 | pRS316-ATG2(T114A)-FLAG | This study | |
| pTKO753 | pRS316-ATG2(S109AS116A)-FLAG | This study | |
| pTKO788 | pRS316-ATG2(S109A T114A S116A)-FLAG | This study | |

| | | |
|---|---|---|
| pTKO754 | pRS316-ATG2(S109DS116D)-FLAG | This study |
| pTKO789 | pRS316-ATG2(S109D T114D S116D)-FLAG | This study |

Supplementary Table 2. Yeast strains used for this study.

| Name | Genotype | Figures |
|---|---|---|
| BY4741 | *MATa his3D1 leu2D0 met15D0 ura3D0* | Ref 64 |
| W303-1A | *MAT*a *ade2-1 ura3-1 his3-11,15 trp1-1 leu2-3,112 can1-100* | Ref 65 |
| ScTK296 | BY4741, *ATG17-2×mCherry-hphNT1 atg2*D::*natNT2* | 5E |
| ScTK298 | BY4741, *his3*D*1*::*PGK1-EGFP-HIS3* atg2D::natNT2 | 5D, 5I, S4E |
| ScTK1751-2 | BY4741, *atg2*D::*natNT2 atg18*D::*hphNT1* | 1H, S1C |
| ScTK1752-2 | BY4741, *his3*D*1*::*PGK1-EGFP-HIS3* atg2D::natNT2 *atg18*D::*hphNT1* | 1I, 1J |
| ScTK1663 | BY4741, *ATG17-2×mCherry-hphNT1 atg2*D::*natNT2 atg8*D*::kanMX4 pRS303-ATG2-3×HA-EGFP* | S4B |
| ScTK1664 | BY4741, *ATG17-2×mCherry-hphNT1 atg2*D::*natNT2 atg8*D*::kanMX4 pRS303-ATG2(DARCH1)-3×HA-EGFP* | S4B |
| ScTK2498 | W303-1A, *ade2-1*::*ADE2 pRS303-$P_{GPD}$-APE1 leu2-3,112*::*mNeonGreen-ATG8-kanMX4 atg2*D::*hphNT1 vps13*D::*zeoNT3 pRS305-mRFP-APE1 pRS306-ATG2-3×HA-egfp(G67A)* | 5J |
| ScTK2499 | W303-1A, *ade2-1*::*ADE2 pRS303-$P_{GPD}$-APE1 leu2-3,112*::*mNeonGreen-ATG8-kanMX4 atg2*D::*hphNT1 vps13*D::*zeoNT3 pRS305-mRFP-APE1 pRS306-atg2(ARCH1*D*21)-3×HA-egfp(G67A)* | 5J |
| ScTK2512 | W303-1A, *ade2-1*::*ADE2 pRS303-$P_{GPD}$-APE1 leu2-3,112*::*mNeonGreen-ATG8-kanMX4 atg2*D::*hphNT1 vps13*D::*zeoNT3 pRS305-mRFP-APE1 pRS306-atg2(ARCH1*D*19)-3×HA-egfp(G67A)* | 5J |
| ScTK2513 | W303-1A, *ade2-1*::*ADE2 pRS303-$P_{GPD}$-APE1 leu2-3,112*::*mNeonGreen-ATG8-kanMX4 atg2*D::*hphNT1 vps13*D::*zeoNT3 pRS305-mRFP-APE1 pRS306-atg2(ARCH1*D*18)-3×HA-egfp(G67A)* | 5J |
| ScTK1702 | W303-1A, *ade2-1*::*ADE2 APE1-mTagBFP2-kanMX4 pRS304-$P_{GPD}$-APE1 pRS306-$P_{GPD}$-APE1 leu2-3,112*::*2×mCherry-ATG8-hphNT1 atg2*D::*natNT2 pRS303-ATG2-3×HA-EGFP* | S4F |
| ScTK1703 | W303-1A, *ade2-1*::*ADE2 APE1-mTagBFP2-kanMX4 pRS304-$P_{GPD}$-APE1 pRS306-$P_{GPD}$-APE1 leu2-3,112*::*2×mCherry-ATG8-hphNT1 atg2*D::*natNT2 pRS303-atg2(ARCH1*D*21)-3×HA-EGFP* | S4F |
| ScTK1704 | W303-1A, *ade2-1*::*ADE2 APE1-mTagBFP2-kanMX4 pRS304-$P_{GPD}$-APE1 pRS306-$P_{GPD}$-APE1 leu2-3,112*::*2×mCherry-ATG8-hphNT1 atg2*D::*natNT2 pRS303-atg2(ARCH1*D*19)-3×HA-EGFP* | S4F |
| ScTK1705 | W303-1A, *ade2-1*::*ADE2 APE1-mTagBFP2-kanMX4 pRS304-$P_{GPD}$-APE1 pRS306-$P_{GPD}$-APE1 leu2-3,112*::*2×mCherry-ATG8-hphNT1 atg2*D::*natNT2 pRS303-atg2(ARCH1*D*18)-3×HA-EGFP* | S4F |

| | | |
|---|---|---|
| SEY6210 | *MATα lys2 suc2 his3 leu2 trp1 ura3* | Ref 67 |
| YOC5209 | SEY6210 *mNeonGreen-ATG8::LEU2* | 6A, S5B, S5C |
| YHL10 | SEY6210, *atg2Δ mNeonGreen-ATG8::LEU2* | 6A |
| YHL11 | SEY6210, *atg2*Δ366-374 *mNeonGreen-ATG8::LEU2* | 6A |
| YHL12 | SEY6210, *atg2*Δ356-374 *mNeonGreen-ATG8::LEU2* | 6A |
| YHL13 | SEY6210, *atg2*Δ354-374 *mNeonGreen-ATG8::LEU2* | 6A |
| ScTK2062 | W303-1A, *ade2-1*::*ADE2 PGK1-EGFP-kanMX4 atg2*Δ::*hphNT1* | 3G, 3H |
| ScTK2514 | W303-1A, *ade2-1*::*ADE2 atg2*Δ::*natNT2 hphNT1-ADHp-yeGFP-MTC4* | 3I |
| ScTK2316 | W303-1A, *ade2-1*::*ADE2 SCS2-6×HA-zeoNT3* | 3J |
| ScTK2317 | W303-1A, *ade2-1*::*ADE2 SCS2-6×HA-zeoNT3 ATG2-3×FLAG-hphNT1* | 3J |
| ScTK2323 | W303-1A, *ade2-1*::*ADE2 SCS2-6×HA-zeoNT3 ATG2-3×FLAG-hphNT1 atg1*Δ::*natNT2* | 3J |
| ScTK2324 | W303-1A, *ade2-1*::*ADE2 SCS2-6×HA-zeoNT3 ATG2-3×FLAG-hphNT1 atg9*Δ::*natNT2* | 3J |
