## Supplementary Data 2 for "Atg1 phosphorylates Atg2 to gate phospholipid flow through a continuous conduit for autophagosome biogenesis"

**ImageJ Script S1**

```
// Select the image at the point where the isolation membrane is most expanded
// Crop and isolate the desired region

// Clear the ROI Manager and Results
if (roiManager("count") > 0) {
	roiManager("reset");
}
run("Clear Results");

//Binarize the image
run("Smooth", "stack");
setAutoThreshold("Otsu dark");
run("Convert to Mask", "method=Otsu background=Dark black");

//Measure the area of the isolation membrane
run("Analyze Particles...", "display exclude add composite stack");
```

**ImageJ Script S2**

```
rename("Atg8");

run("Smooth");
run("Sharpen");

//maximum intensity projection
run("Z Project...", "projection=[Max Intensity]");
run("Split Channels");

//extract Ape1 complex
selectWindow("C2-MAX_Atg8");
setAutoThreshold("Otsu dark");
//run("Threshold...");
setOption("BlackBackground", true);
```

```
run("Convert to Mask");
run("Maximum...", "radius=2");

//find the isolation membranes
selectWindow("C1-MAX_Atg8");
setAutoThreshold("Moments dark");
run("Convert to Mask");
imageCalculator("Multiply create", "C1-MAX_Atg8","C2-MAX_Atg8");
selectWindow("Result of C1-MAX_Atg8");
run("Analyze Particles...", "size=0.01-Infinity exclude add in_situ");

//Measure the perimeter of the isolation membrane
roiManager("Measure");

roiManager("Deselect");
roiManager("Delete");
run("Close All");
```

**ImageJ Script S3**

```
rename("2G17C");
//maximum intensity projection
run("Z Project...", "projection=[Max Intensity]");
run("Split Channels");

//extract Atg17 puncta
selectWindow("C1-MAX_2G17C");
run("Find Maxima...", "prominence=1000 output=[Single Points]");
run("Maximum...", "radius=3");

//extract Atg2 puncta
selectWindow("C2-MAX_2G17C");
run("Find Maxima...", "prominence=500 output=[Single Points]");
run("Maximum...", "radius=3");
```

```
//extract colocalized puncta
imageCalculator("Multiply create", "C1-MAX_2G17C Maxima","C2-MAX_2G17C
Maxima");
selectWindow("Result of C1-MAX_2G17C Maxima");
run("Find Maxima...", "prominence=1 output=Count");
selectWindow("C1-MAX_2G17C Maxima");
run("Find Maxima...", "prominence=1 output=Count");
run("Close All");;
```
